# A unique H2B acetylation signature marks active enhancers and predicts their target genes

**DOI:** 10.1101/2022.07.18.500459

**Authors:** Takeo Narita, Yoshiki Higashijima, Sinan Kilic, Tim Liebner, Jonas Walter, Chunaram Choudhary

**Author notes:** These authors contributed equally to this work (Takeo Narita, Yoshiki Higashijima).

## Abstract

Chromatin features are widely used for genome-scale mapping of enhancers. However, discriminating active enhancers from other cis-regulatory elements, predicting enhancer strength, and identifying their target genes remains challenging. Here we establish histone H2B N-terminus multisite lysine acetylation (H2BNTac) as a genuine signature of active enhancers. H2BNTac prominently marks candidate active enhancers and their target promoters and discriminates them from ubiquitously active promoters. Two mechanisms afford the distinct H2BNTac specificity. (1) Unlike H3K27ac, H2BNTac is specifically catalyzed by CBP/p300. (2) H2A-H2B, but not H3-H4, are rapidly exchanged through transcription-induced nucleosome remodeling. H2BNTac-positive candidate enhancers show a high validation rate in orthogonal enhancer activity assays, and a vast majority of endogenously active enhancers are marked by H2BNTac and H3K27ac. Notably, H2BNTac intensity predicts enhancer strength and outperforms the current state-of-the-art models in predicting enhancer target genes. These findings have broad implications for generating fine-grained enhancer maps and modeling enhancer-dependent gene regulation.

## Introduction

Cis-regulatory enhancers are instrumental for activating signal-induced and developmentally regulated genes. Candidate enhancers are identified using massively parallel reporter assay (MPRA), enhancer RNA (eRNA) transcription, and enrichment of certain chromatin features^1-3^. Chromatin features, such as H3K4me1, H3K27ac, and DNase I hypersensitivity (DHS), are widely used for genome-scale enhancer mapping. A drawback of DHS is that it merely signifies open chromatin without discriminating enhancers and promoters, and the downside of H3K4me1 is that it poorly discriminates between active and poised enhancers. Compared to other chromatin marks, H3K27ac shows greater enhancer specificity. For this reason, H3K27ac is widely used, and it is the only acetylation mark that is broadly profiled by the International Human Epigenome Consortium partners, including ENCODE, BLUEPRINT, and the NIH Epigenome Roadmap^4-6^.

While H3K27ac and other chromatin features have tremendously facilitated genome-scale mapping of candidate enhancers, some challenges still remain that hamper a deeper understanding of enhancer-dependent gene regulation. (1) It is often discussed that distal enhancers encompass heterogeneous groups and thousands of them apparently lack H3K27ac^7-10^. However, extent of the heterogeneity and fraction of active mammalian enhancers that lack H3K27ac remains unclear. (2) H3K27ac poorly discriminates between proximally occurring active enhancers and promoters^3,11,12^. H3K4me1/H3K4me3 ratio can discriminate candidate enhancers from promoters, but poorly discriminate active regions from poised and inactive regions. (3) Enhancers predicted by chromatin marks show variable validation rates in orthogonal assays, giving the notion that chromatin marks are poor predictors of endogenously active enhancers^13-15^. (4) A key goal of mapping enhancers is to estimate their functional impact on gene activation. However, despite the availability of many high-quality epigenome reference datasets, predicting native enhancer strength and enhancer target genes remain challenging^16,17^.

This work was motivated by our prior findings showing that (1) in addition to H3K27ac, CBP/p300 catalyzes H2B N-terminus multisite lysine acetylation (H2BNTac) with rapid kinetics^18^, (2) lysine deacetylase (KDAC) inhibition strongly increases H2BNTac^19^, (3), some of the H2BNTac sites occur at a relatively high stoichiometry^20^, and (4), kinetic CBP/p300 activity controls enhancer-mediated transcription activation^21^. What remained unclear was the relative specificity of H3K27ac and H2BNTac sites in marking candidate enhancers, the role of CBP/p300 in locus-specific regulation of these marks, and the usefulness of H2BNTac in predicting enhancer strength and target genes. Through systematic comparison of H3K27ac and H2BNTac, here we show that genomic occupancy of H2BNTac sites is distinct from H3K27ac and other frequently analyzed chromatin marks. H2BNTac sites distinctively mark active enhancers and discriminate them from other candidate cis-regulatory elements (cCREs). H2BNTac marked regions show a high validation rate in orthogonal enhancer activity assay. Importantly, H2BNTac most accurately defines locus-specific CBP/p300 activity and outperforms H3K27ac in predicting enhancer target genes and the degree of their enhancer dependency.

## Results

### Multiple H2BNTac sites occupy the same genomic regions

In proteomic analyses, H3K27ac and H2BNTac sites are similarly regulated by CBP/p300^18^ (**Extended Data Fig. 1**), yet the genome occupancy patterns of H2BNTac sites are dissimilar from each other and variable across cell types^22,23^ (**Supplemental Note 1**). The reasons for this discordance have remained unclear. As it stands, it is not known if CBP/p300 similarly or differentially regulates H3K27ac and H2BNTac at individual loci. To resolve this conundrum, we systematically investigated H3K27ac and H2BNTac genomic occupancy and regulation by chromatin immunoprecipitation sequencing (ChIP-seq). To allow reproducibility, all ChIP-seq analyses were performed using rabbit monoclonal H2BNTac antibodies. We tested all H2BNTac antibodies that were commercially available at the start of this work, and found six, targeting H2BK5ac, H2BK12ac, H2BK16ac, and H2BK20ac, that passed quality control in ChIP-PCR analyses (**Supplementary Table 1**). For clarity, antibodies targeting the same sites are distinguished by their clone’s name as H2BK5ac^EP857Y^, H2BK5ac^D5H1S^, H2BK20ac^EPR859^, and H2BK20ac^D7O9W^.

H2BNTac occupancy and regulation were analyzed in mouse embryonic stem cells (mESC) treated with or without the CBP/p300 catalytic inhibitor A-485^24^. A roughly similar number of peaks were identified for H3K27ac, H2BK5ac^EP857Y^, H2BK16ac, H2BK20ac^EPR859^, and H2BK20ac^D7O9W^, but fewer peaks were identified for H2BK5ac^D5H1S^ and H2BK12ac (**Extended Data Fig. 2**). This difference was not due to a lower sequencing depth, but due to lower H2BK5ac^D5H1S^ and H2BK12ac peak intensity (**Extended Data Fig. 2**), which possibly reflects weaker enrichment efficiency of the antibodies and/or lower abundance of these marks. Regardless, a majority of H2BNTac site peaks overlapped with each other, and 92-99% of H2BK5ac^D5H1S^ and H2BK12ac peaks overlapped with H2BK16ac and H2BK20ac (**Extended Data Fig. 3a**). The peak height of overlapping H2BNTac peaks was significantly greater than non-overlapping peaks. As expected, most (71-92%) H3K27ac and H2BNTac peaks occurred in ATAC (Assay for Transposase-Accessible Chromatin) accessible regions, and non-overlapping peaks were low abundant (**Extended Data Fig. 3b**). These results show that most H2BNTac sites occur in the same, openly accessible regions.

### H2BNTac partially overlaps with H3K27ac

We analyzed the overlap of H3K27ac with H2BNTac as well as H3K9ac, which is primarily acetylated by GCN5/PCAF. 91% of H3K27ac peaks overlapped with H2BK5ac^EP857Y^, 72% overlapped with H3K9ac, and 38-62% with other H2BNTac sites (**Fig. 1a**). In comparison to H3K27ac overlapping peaks, the height of non-overlapping H2BNTac peaks was very low. This show that almost all strongly marked H2BNTac^+^ regions are marked with H3K27ac, but many abundantly marked H3K27ac^+^ regions lack H2BNTac.

**Fig 1.**
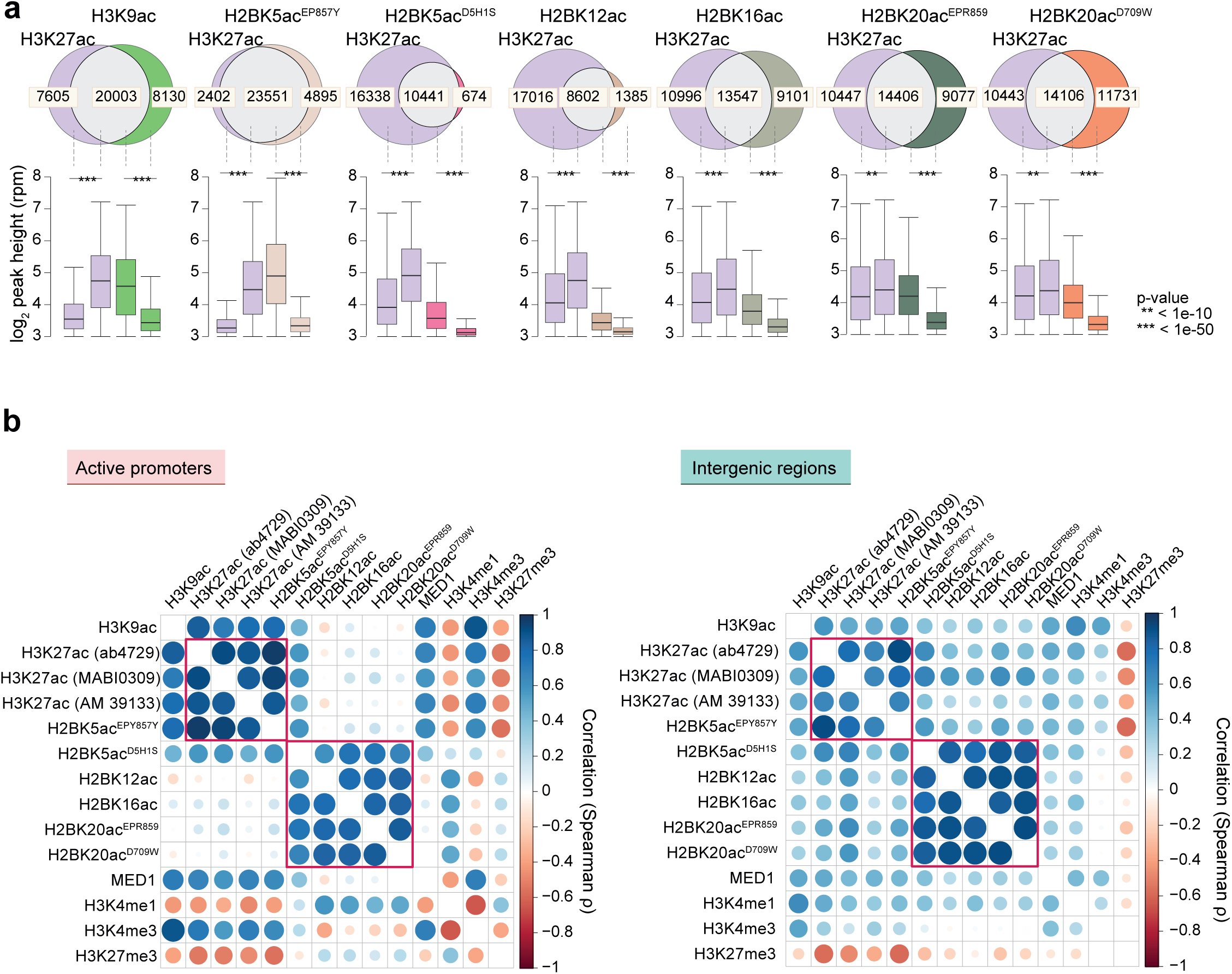
A majority of H2BNTac site peaks overlap with H3K27ac. **a,** Venn diagrams show the overlap between the number of ChIP-seq peaks identified for the indicated histone acetylation marks. Box plots, below each Venn diagram, show the ChIP signal intensity of the overlapping and non-overlapping peaks. One H2BNTac peak can overlap with more than one H3K27ac peak or versa, vice which leads to slight differences in the total number of H3K27ac peak counts in different Venn diagrams. The box plots display the median, upper and lower quartiles, and whiskers show 1.5× interquartile range (IQR). **b,** Correlation among H2BNTac sites and the other indicated chromatin marks. Pair-wise correlation (Spearman ρ) was determined using the normalized ChIP-seq counts. The left panel shows a correlation at active promoter regions (±1 kb from transcriptional start sites), and the right panel shows correlations at intergenic regions. Color intensities and the size of the circles indicate correlation (Spearman ρ).

### H2BNTac distinctly correlates with other chromatin marks

To evaluate H2BNTac enrichment in different regulatory regions, we analyzed H2BNTac correlation with chromatin features associated with promoters (H3K4me3, H3K9ac), enhancers (H3K4me1), both at promoters and enhancers (MED1, H3K27ac), or Polycomb-repressed regions (H3K27me3). As expected, H3K4me3 positively correlated with H3K9ac and MED1, and H3K27ac but negatively correlated with H3K27me3 (**Extended Data Fig. 4a**). H2BK5ac^D5H1S^, H2BK12ac, H2BK16ac, and H2BK20ac strongly correlated with each other (Spearman ρ: 0.84-0.93; Pearsońs r: 0.85-0.94) but modestly correlated (Spearman ρ: 0.30-0.72; Pearsońs r: 0.24-0.73) with H3K27ac, H3K9ac, and MED1, and weakly or negatively correlated (Spearman ρ: −0.55-0.20; Pearsońs r: −0.53-0.15) with H3K4me3 and H3K27me3. H2BK5ac^EP857Y^ correlated most strongly with H3K27ac. H3K27me3 is more strongly negatively correlated with H3K27ac than H2BNTac, likely because of the mutual exclusivity of H3K27ac and H3K27me3. We separately analyzed correlations at active promoters and intergenic regions. In intergenic regions, H2BNTac positively correlated with H3K27ac, H3K9ac, and MED1, but at active promoters, H2BNTac showed little correlation with any of these marks (**Fig. 1b**). After completing all the other experiments and analyses, we obtained an additional antibody recognizing H2BK11ac. ChIP-seq analyses show that H2BK11ac peaks extensively overlap (84-96%) with other H2BNTac sites, and H2BK11ac shows a similar occupancy profile as other H2BNTac sites (**Extended Data Fig. 4b-c**).

The H2BK5ac^EP857Y^ occupancy profile was a notable outliner among the H2BNTac sites (**Fig. 1b**). Antibody specificity is often a concern^25^. We noted a high sequence similarity near H2BK5 and H3K27, and H2BK20 and H2BK120. Therefore, we assessed the specificity of H2BK5ac, H2BK20ac, and H3K27ac antibodies used in our work, as well as previously used H2BK5ac and H2BK120ac antibodies^22,23^ (**Extended Data Fig. 5a-e**). H2BK5ac^EP857Y^ and H2BK5ac^polyclonal^ antibodies cross-reacted with acetylated H3 peptide, and both H2BK120ac^polyclonal^ antibodies cross-reacted with H2BNTac, but H2BK20ac antibodies did not cross-react with H2BK120ac. H2BK5ac^D5H1S^ did not show measurable cross-reactivity in quantitative image-based cytometry, but in ChIP-seq, it showed a higher correlation with H3K27ac than other H2BNTac sites antibodies (**Fig. 1b**), indicating that H2BK5ac^D5H1S^ may weakly recognize H3K27ac. These cross-reactivities can rationalize unexpected and strong correlation between the ChIP-seq profiles of H2BK5ac^EP857Y^ and H3K27ac (**Fig. 1b**) and between H2BK20ac and H2BK120ac^22,23^. Because of the cross-reactivity, H2BK5ac^EP857Y^ was excluded from defining the H2BNTac signature.

### H2BNTac proficiently marks distal enhancers

Just as H3K27ac, MED1 binding also marks candidate enhancers^26,27^. We determined H2BNTac occupancy at MED1^+^ intergenic regions. Acetylation site peaks were grouped into quartiles, and the fraction of peaks mapping to MED1^+^ intergenic regions was determined. H2BK16ac, H2BK20ac, ad H3K27ac marked a majority of MED1^+^ intergenic regions (**Fig. 2a**). A smaller fraction of MED1^+^ regions were marked by H2BK12ac and H2BK5ac^D5H1S^, reflecting limited coverage of these marks in ChIP-seq (**Extended Data Fig. 2**). H3K27ac and H2BNTac almost completely overlapped with MED1^+^ super-enhancer regions^27^ (**Fig. 2b**). These results show a similarly high sensitivity of H3K27ac and H2BNTac in detecting MED1^+^ candidate enhancers.

**Fig. 2.**
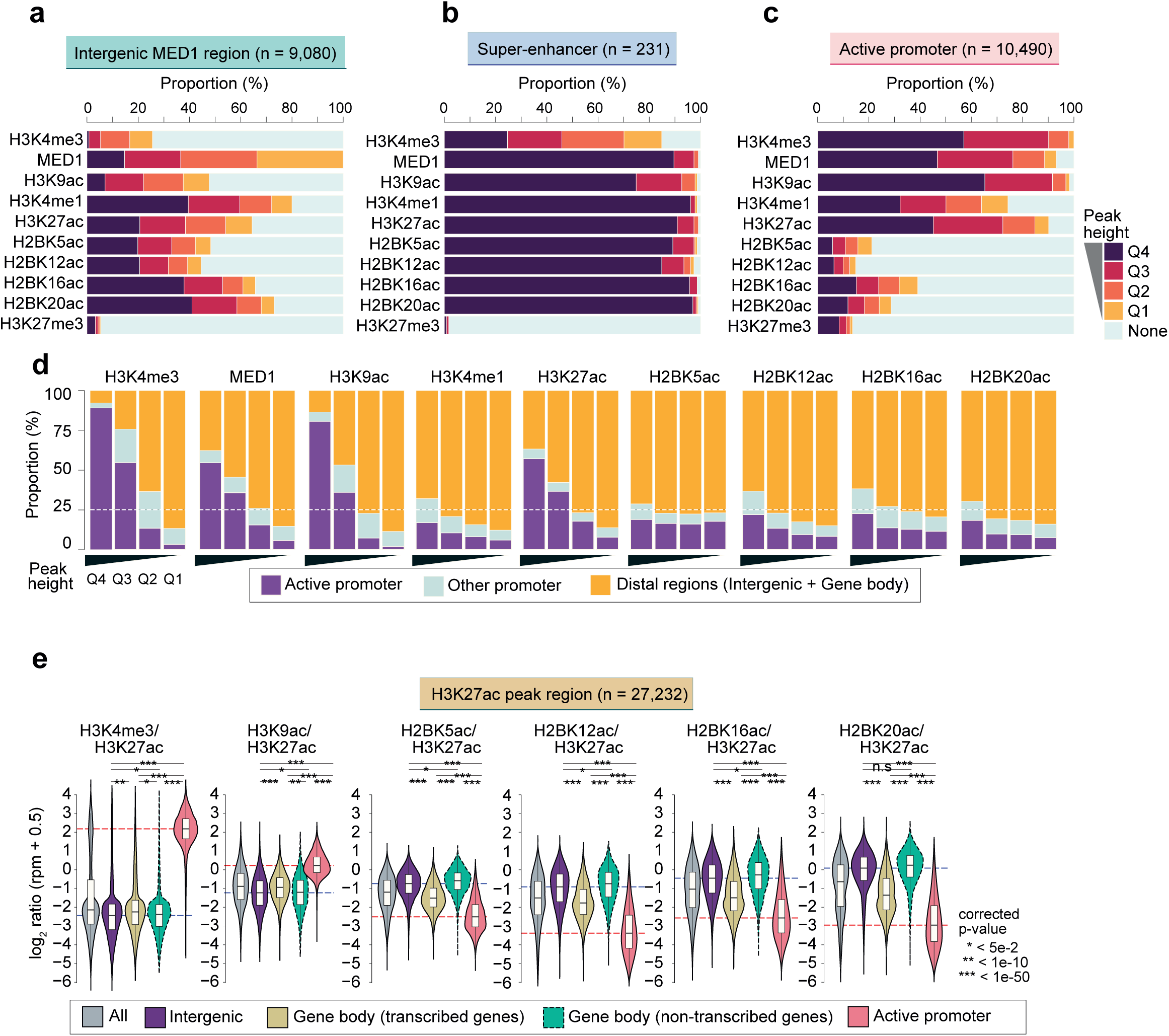
H2BNTac prominently marks enhancers and discriminates them from active promoters. **a-c,** H2BNTac sites more prominently mark enhancers than promoters. Based on peak height, the indicated chromatin marks are grouped into quartiles (Q4-Q1). Shown is the fraction of the indicated chromatin mark peaks mapping to MED1 occupied intergenic regions (a), MED1 occupied super-enhancers (b), and active promoters (c). Active promoters (±1 kb from TSS) are defined as those mapping to genes expressed in mESC (EU-seq TPM ≥2) and marked with H3K4me3. **d,** Shown is the fraction of the indicated chromatin marks mapping to the depicted genomic regions. For each chromatin mark, peaks were grouped into quartiles (Q4-Q1), and in each category, the fraction of peaks mapping to the indicated genomic regions is depicted. Active promoters are defined as in (a) and the remaining gene TSS are classified as Other promoters. **e,** H2BNTac signal is higher in distal candidate enhancers than in active promoters. Shows are the relative ChIP signal intensities of H3K27ac and H2BNTac sites in H3K27ac-occupied regions. H3K27ac peak regions are grouped into the following categories: All, all peaks; Intergenic, peaks occurring outside promoters and gene bodies; gene body (transcribed genes, TPM ≥2), peaks occurring within actively transcribed gene body; gene body (non-transcribed genes), peaks occurring within the non-transcribed gene body. Active promoters were defined as in (a). The dotted lines indicate the ratio at intergenic median entergenic and active promoter regions. Statistical significance determined by Mann–Whitney U test.

The transcription factors NANOG and OCT4 are enriched in mESC enhancers^28^. NANOG/OCT4 occupied a larger fraction of H2BNTac^+^ regions than H3K27ac^+^ regions (**Extended Data Fig. 6**). Notably, in H2BNTac^+^ regions, NANOG/OCT4 binding was similar in promoter and distal regions, whereas in H3K27ac^+^ and MED1^+^ regions, NANOG/OCT4 binding was much lower in promoters than in distal regions. In the top quartile of H2BNTac^+^ regions, 88-97% were bound by NANOG/OCT4.

### H2BNTac poorly marks active promoters

In mESC, a majority (90-98%) of promoters were marked by H3K27ac, MED1, and H3K9ac, but H2BNTac was not detectable in a large portion of promoters (**Fig. 2c**). In the top quartile, most H3K4me3, H3K9ac, H3K27ac, and MED1 peaks occurred in promoters, whereas most H3K4me1 and H2BNTac peaks occurred in distal regions (**Fig. 2d**). The relative signal of chromatin marks also differed greatly. In H3K27ac^+^ regions, a high H3K9ac/H3K27ac ratio discriminated promoters from distal regions, and a high H3K4me3/H3K27ac ratio even more prominently separated active promoters from distal regions (**Fig. 2e**). Notably, the H2BNTac/H3K27ac ratio showed an opposite pattern, and a high level of H2BNTac positively separated H3K27ac^+^ distal regions from active promoters. Analysis of H3K9ac^+^ regions and H2BK20ac^+^ regions further confirmed these differences (**Extended Data Fig. 7a, b**).

### H2BNTac abundance differs in intergenic and transcribed gene body regions

We used A-485^24^ to investigate locus-specific regulation of H2BNTac and H3K27ac by CBP/p300. A-485 more strongly downregulated H2BNTac in active promoters and actively transcribed gene body regions than in distal regions (**Fig. 3a**, **Extended Data Fig. 8a-b**). H2BNTac is increased by class I deacetylases inhibitors in vivo and HDAC1/2 can deacetylate it in vitro ^19,29^. We tagged endogenous HDAC1/2 with GFP-FKBP^12F36V^, depleted HDAC1/2- GFP-FKBP^12F36V^ using the dTAG approach^30^, and confirmed that H2BNTac is deacetylated by HDAC1/2 (**Extended Data Fig. 9**). Together, these analyses show that most H2BNTac is catalyzed by CBP/p300 and H2BNTac is more rapidly turned over in actively transcribed regions than in intergenic and non-transcribed genic regions.

**Fig. 3.**
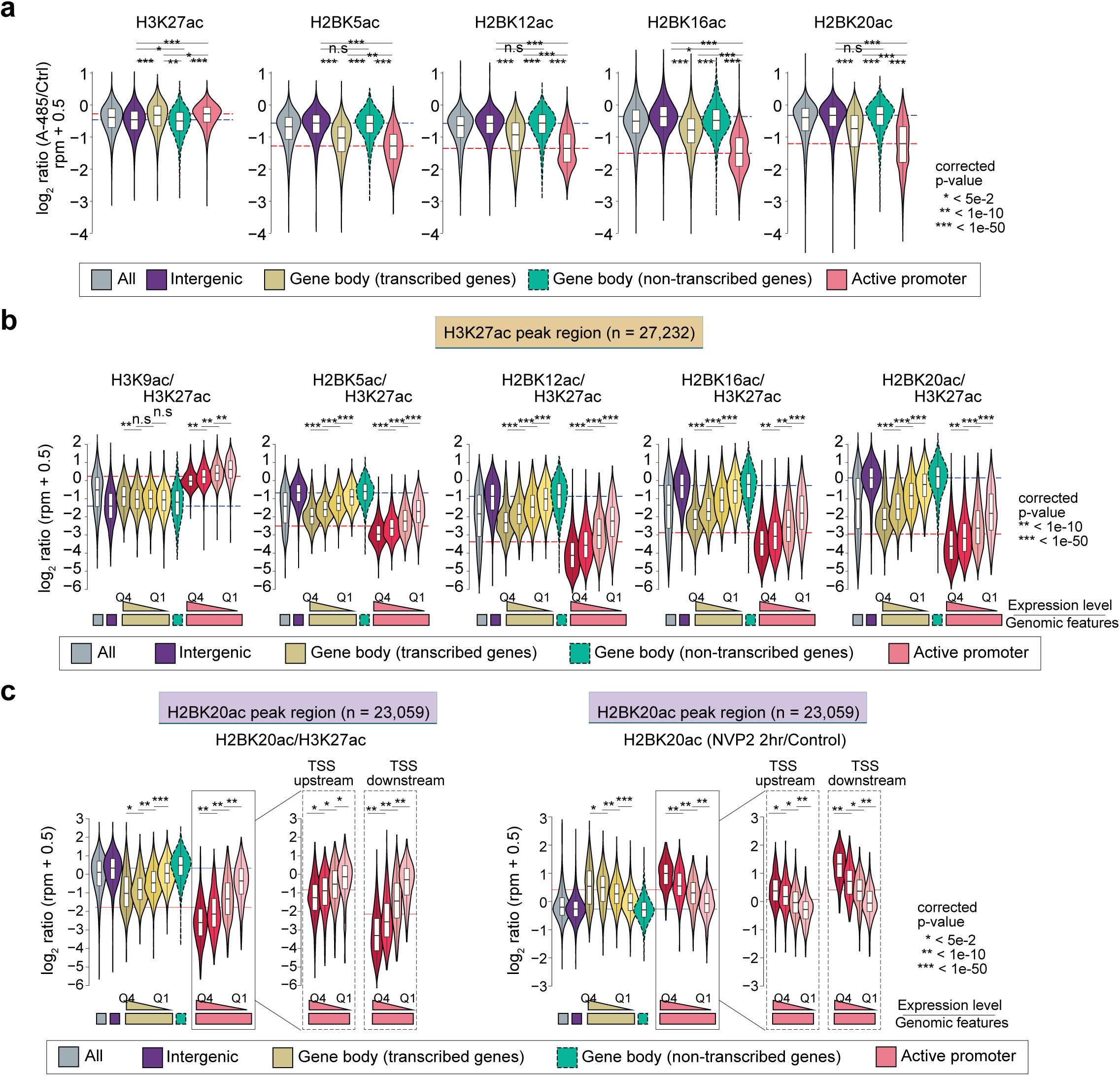
CBP/p300 regulates global H2BNTac, and the H2BNTac signal is lower in actively transcribed regions. **a,** CBP/p300 inhibition leads to a more rapid decline of H2BNTac in actively transcribed regions than in intergenic regions. Shown is the relative H3K27ac and H2BNTac site ChIP signal intensity in untreated and A-485 treated (15 min) cells. Different genomic regions were classified as defined in Fig. 1e. Dotted lines indicate the median ratio at intergenic and active promoter regions. **b,** H2BNTac signal is lower in highly transcribed regions. Shown is the ratio of the indicated histone marks in H3K27ac marked regions. Different genomic regions were classified as defined in Fig. 1e. Based on the nascent transcription levels the corresponding genes, gene body and active promoter regions were further subdivided into quartiles. **c,** Transcription inhibition increases H2B acetylation in actively transcribed regions. Transcription was inhibited using the CDK9 inhibitor NVP-2 (2 hours), and H2BK20ac was analyzed in control and NVP-2 treated cells using ChIP-seq. Shown is the relative ratio of H2BK20ac/H3K27ac in untreated conditions (left, panels), and the relative change in H2BK20ac in NVP2 treated cells (right panels). Different genomic regions were classified as defind in Fig. 2e. Based on the nascent transcription levels the corresponding genes, gene body and active promoter regions were further subdivided into quartiles. Moreover, in active promoters, TSS upstream (0--2kb from TSS) and TSS downstream (0- +2kb from TSS) regions were analyzed separately. Statistical significance determined by Mann–Whitney U test.

### Transcription-coupled nucleosome exchange impacts H2BNTac genomic occupancy

We were intrigued to find that the H2BNTac/H3K27ac and H2BNTac/H3K9ac ratio is lower in the actively transcribed gene body regions than in intergenic and non-transcribed gene body regions (**Fig. 2e**). To explore this further, we grouped genes based on their nascent transcript expression levels. The H2BNTac/H3K27ac ratio was inversely associated with gene transcription; the ratio was lowest in genes with the highest expression (**Fig. 3b, Extended Data Fig. 10a-b**). To investigate the link between active transcription and H2BNTac, transcription was inhibited using the CDK9 inhibitor NVP2 (2h, 100nM)^31^, and the change in H2BK20ac was analyzed by ChIP-seq. The H2BK20ac/H3K27ac ratio was more strongly increased by NVP2 in actively transcribed regions than in non-transcribed regions (**Fig. 3c, Extended Data Fig. 10c**). The increase in H2BK20ac directly relates to gene expression levels; H2BK20ac is most strongly increased in highly transcribed regions. In promoters, the H2BK20ac/H3K27ac ratio is different between TSS upstream and downstream regions. After NVP2 treatment, H2BK20ac is more strongly increased in TSS downstream than in upstream regions (**Fig. 3c**). This shows that ongoing transcription causes loss of H2BNTac. In retrospect, this result can be rationalized by the transcription-induced selective exchange of H2A-H2B dimer(s), and redeposition of H3-H4 tetramer^32,33^.

### Enhancer specificity of H2BNTac in human cells

To confirm H2BNTac specificity in human cells, we used H2BK20ac as a representative H2BNTac mark and performed ChIP-seq analyses in K562, a chronic myelogenous leukemia cell line that is included among ENCODÉs Tier 1 cell lines. The relative H2BK20ac, H3K27ac, H3K4me3, and H3K9ac enrichment were analyzed in active promoters and distal regions. Similar to mESC, in H3K27ac^+^ regions, the H2BK20ac/H3K27ac ratio was much higher in distal regions than in active promoters (**Fig. 4a**). The H2BK20ac/H3K27ac ratio provided much-improved discrimination between distal enhancers and promoters than the H3K27ac/H3K9ac ratio. The H2BK20ac/H3K27ac ratio in actively transcribed gene body regions was lower than in distal regions. The H2BK20ac/H3K27ac ratio in distal enhancers and active promoters is the opposite of the H3K4me3/H3K27ac ratio. Similarly, in H3K9ac^+^ regions, the H3K27ac/H3K9ac ratio was only modestly higher at distal candidate enhancers than promoters, whereas H2BK20ac/H3K9ac ratio was much higher in enhancers (**Fig. 4a**). In ChromHMM-defined states in mESC and K562^34^, H2BNTac was less prominently enriched at promoters than H3K27ac but was enriched at a comparable or higher level in candidate enhancer regions (**Extended data Fig. 11, 12**).

**Fig. 4.**
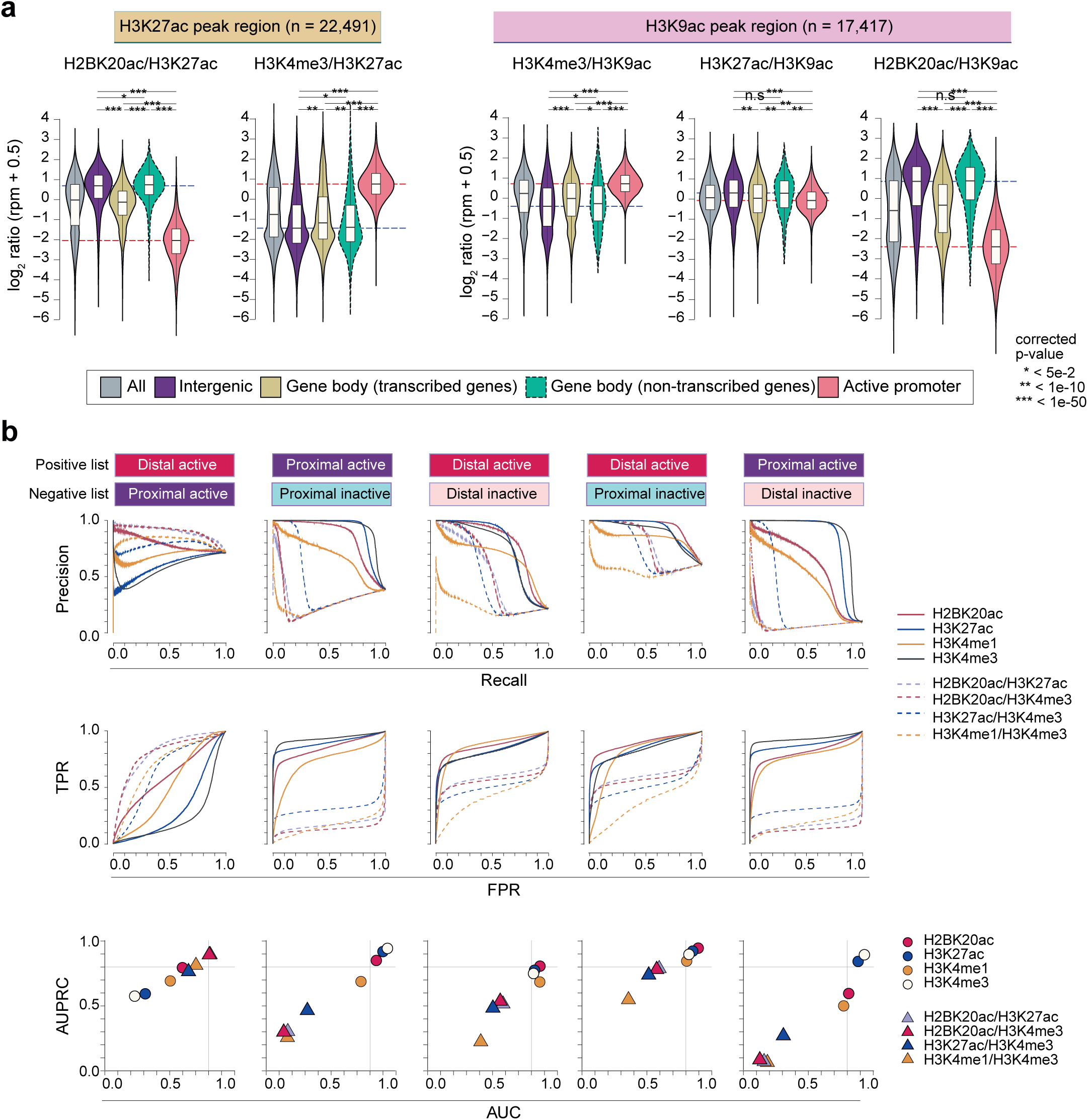
H2BNTac segregates enhancers and active gene promoters in human cells. **a,** Shown are the relative ChIP-seq signal intensities of the indicated histone marks at H3K27ac^+^ (left) or H3K9ac^+^ (right) genomic regions. H3K27ac and H3K9ac peak regions were further grouped into the indicated categories, as specified in Fig. 1e. Active promoters are defined as TSS-proximal regionsns (±1 kb from TSS) of genes that are actively transcribed in K562 (RNA-seq, TPM ≥2) and whose promoters are marked with H3K4me3. Statistical significance determined by Mann–Whitney U test. **b,** Combination of H2BNTac and other histone marks discriminate transcriptionally active and inactive PINTS regions. Transcriptionally active and inactive regions in K562 were defined based on RNA expression evidence in K562 and non-K562 cell lines^14^. PINTS regions were further classified as proximal or distal as defined by Yao et.al. ^14^ (see Methods). The discriminatory potential of different histone marks was analyzed by comparing ChIP-seq signal enrichment within +/-500bp of PINTS region centers. Receiver operating characteristics (ROC) and precision-recall curves (PRC) were generated under the different thresholds of peak enrichment or peak enrichment ratios.

Manual inspection of several loci confirmed differential H2BNTac and H3K27ac occupancy. H3K27ac contiguously marks promoters and upstream regions in *Eif4a2, Actb*, and *Taf1*, whereas H2BNTac only marked the upstream regions near *Eif4a2 and Actb* (**Extended Data Fig. 13a**). The *Taf1* upstream region lacked H2BNTac, had high H3K4me3, and high nascent transcription, indicating that *Taf1* upstream H3K27ac peak denotes an unannotated alternative promoter. H2BNTac is also poorly detectable in promoters of several snoRNA genes, which, unlike canonical promoters, lack the promoter marker H3K4me3 (**Extended Data Fig. 13b**), have previously been mistakenly interpreted as candidate enhancers^27,35^, possibly because of the lack of H3K4me3. Differential H3K27ac and H2BNTac occupancy was also confirmed in several human loci (**Extended Data Fig. 14**).

### H2BNTac complements other histone modifications in discriminating CREs

We investigated if H2BNTac can be combined with other chromatin features to discriminate between active and inactive CREs. As a reference, we used transcriptionally active and inactive regions defined by PINTS (peak identifier for nascent transcript starts)^14^. PINTS regions were classified into four categories. (1) proximal active (active PINTS regions in K562, localizing within 500bp from TSS), (2) proximal inactive (inactive PINTS regions in K562, localizing within 500bp from TSS), (3) distal active (active PINTS regions in K562, localizing >500bp away from TSS), and (4) distal inactive (distal PINTS regions that are inactive in K562 and occur >5kb away from TSS and >2kb away from any distal active regions in this cell line) (see Methods for further details). H3K4me3 was best at separating proximal active from inactive regions (AUPRC: 0.94; AUROC: 0.93), while H2BK20ac was best at separating distal active from inactive regions (AUPRC: 0.87; AUROC: 0.81) (**Fig. 4b, Extended Data Fig. 15**). Notably, high H2BK20ac/H3K4me3 and H2BK20ac/H3K27ac ratios outperformed the H3K4me1/H3K4me3 ratio in separating distal active from proximal active regions. The discriminatory potential of these marks is exemplified in the region near *Ahnak* which harbors about two dozen active promoters and a few candidate enhancers (**Extended Data Fig. 16**). Active promoters are strongly marked with H3K4me3 and H3K27ac but display low H2BNTac, whereas H2BNTac prominently marks candidate enhancer regions near *Ahnak*. Of note, both H2BNTac and H3K4me1 show preferential enrichment in distal candidate enhancer regions, but their occupancy profiles are non-identical. Gene synteny in this region is almost fully conserved between mice and humans, and the analogous region in K562 also displays differential occupancy of these marks (**Extended Data Fig. 16**). These results demonstrate that genomic occupancy profiles of H2BNTac, H3K27ac, H3K4me3, and H3K4me1 are non-identical. While none of the marks display absolute specificity, H2BNTac complements other marks and affords valuable information in confidently discerning candidate active enhancers.

### Validation of H2BNTac^+^ candidate enhancers by MPRA

Next, we validated the enhancer activity of H2BNTac^+^ regions by using candidate enhancers identified by massively parallel reporter assay (MPRA)^36^. From 25,643 MPRA^+^ regions, only natively accessible (ATAC^+^) regions (n=10,531) were considered as likely active endogenous enhancers. Overall, 31% of H3K27ac^+^ and 31-53% of H2BNTac^+^ regions overlapped with ATAC^+^MPRA^+^ peaks (**Fig. 5a**). Thus, the validation rate of H2BNTac^+^ regions compares favorably with the validation rate (29%) of eRNA-defined candidate enhancers^14^. The number of ATAC^−^MPRA^+^ regions is greater than ATAC^+^MPRA^+^ regions, but only a small fraction of ATAC^−^MPRA^+^ peaks overlapped with H2BNTac^+^ regions. This indicates that H2BNTac specifically marks endogenously accessible candidate enhancer regions, but MPRA can additionally score candidate enhancers that are inaccessible in the native context.

**Fig. 5.**
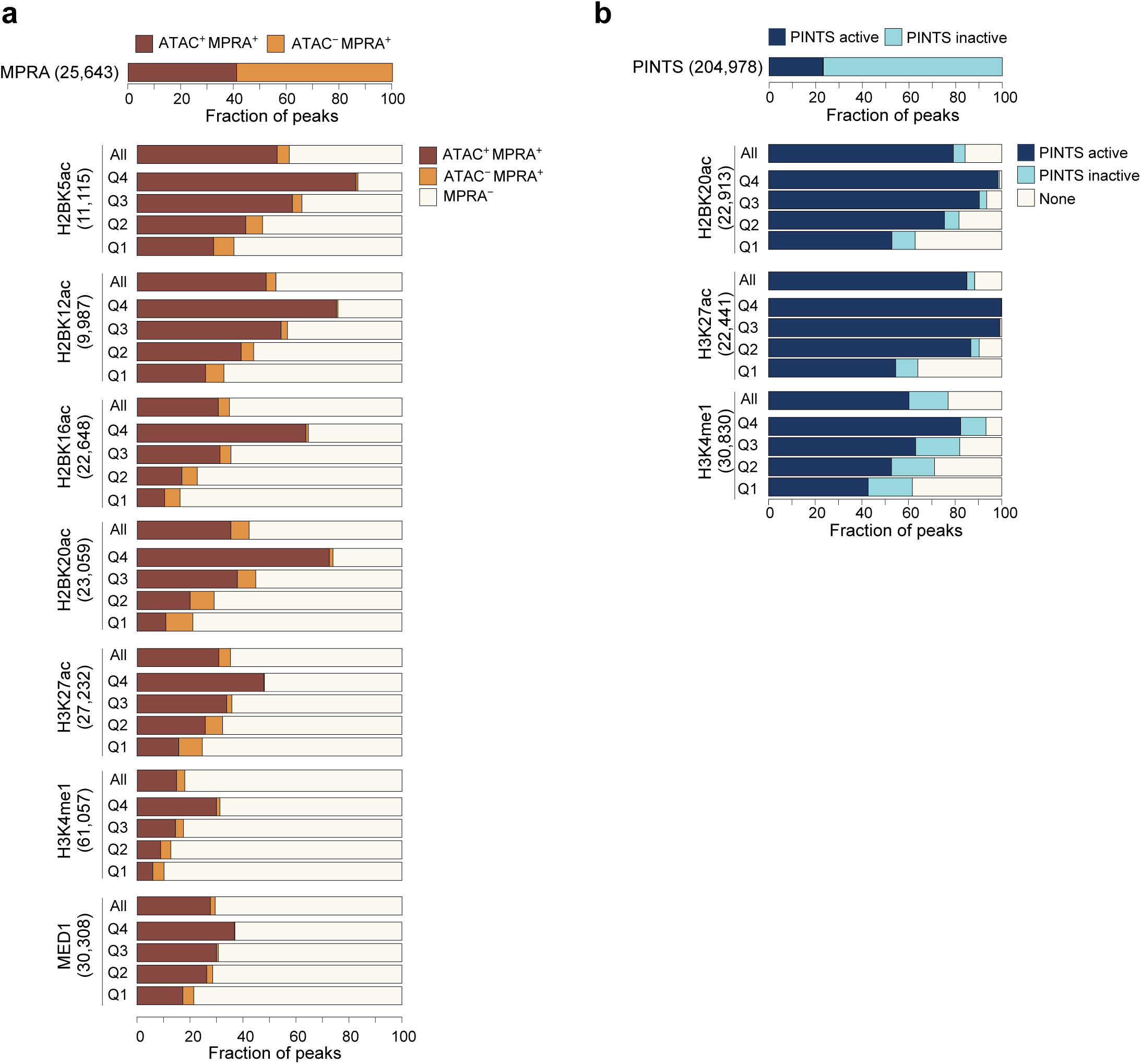
H2BNTac+ regions show a high validation rate in orthogonal assays. **a,** Validation of enhancer activity of H2BNTac+ regions by massively parallel reporter assay (MPRA). Shown are the fraction of H2BNTac, H3K27ac, H3K4me1, and MED1 regions overlapping with ATAC^+^MPRA^+^ or ATAC^−^MPRA^+^ regions in mESC. The top bar chart shows the fraction of MPRA peaks with or without the ATAC signal. The lower bar charts show the fraction of the indicated chromatin mark ChIP seq peaks that overlap with ATAC^+^MPRA^+^ and ATAC^−^MPRA^+^ regions. The indicated chromatin marks were grouped into quartiles (Q4-Q1) based on ChIP-seq signal intensity, and the overlap with MPRA regions is shown within each quartile as well as for all peaks (All). b, A majority of H2BNTac^+^ regions are actively transcribed in vivo. PINTS active regions refer to regions that are actively transcribed in K562, and PINTS inactive regions refer to regions that are not transcribed in K562 but transcribed in other human cells and tissues^14^ (see Methods). The top bar chart shows the fraction of PINTS regions that are active or inactive in K562. The lower bar charts show the fraction of the indicated chromatin mark ChIP seq peaks that overlap with PINTS active and PINTS inactive regions. None: no PINTS classification available for these regions.

Of note, the total number of ATAC^+^MPRA^+^ regions is smaller than the number of H2BNTac^+^ regions, and hence, ATAC^+^MPRA^+^ is not expected to validate all H2BNTac^+^ regions. The MPRA dataset only covered 83% of the genome (with ∼9X coverage)^36^. Assuming that the input library was not biased by the enhancer regions, the maximum expected validation rate in any quartile would be ∼83%. Remarkably, in the top quartile of H2BNTac^+^ regions, 61-82% of H2BNTac regions overlapped with ATAC^+^MPRA^+^ regions (**Fig. 5a**), indicating that the validation rate in the top quartile reaches near saturation.

### Validation of H2BNTac^+^ candidate enhancers by eRNA expression

eRNA transcription is another reliable surrogate of endogenously active enhancers^13,14^. A recent study cataloged hundreds of thousands of endogenously active transcript start sites (TSSs), representing the TSS of annotated genes and candidate enhancers^14^. We generated a reference dataset by combining PINTS-identified nascent TSS sites in >110 cell types and used this large compendium to systematically validate transcription activity in H2BNTac^+^ regions. Of the reference PINTS regions, ∼50,000 were active in K562, and the remaining were active in other cell types or tissues but inactive in K562. Active TSS in K562 are considered “active PINTS”, and TSS that are inactive in K562 are considered “inactive PINTS.” Using the original classification^14^, PINTS regions were classified as proximal (TSS of annotated genes) or distal (distal TSS, not assigned to annotated genes). Overall, ∼79-85% of H2BK20ac and H3K27ac regions overlapped with active PINTS regions (**Fig. 5b**). From this, we estimate the positivity rate to be ∼80% for H3K27ac and/or H2BK20ac regions for endogenous RNA transcription. Among the top 25% H3K27ac^+^ and H2BK20ac^+^ regions, virtually all overlapped with active PINTS regions, showing that almost all abundantly marked H2BNTac and/or H3K27ac regions are actively transcribed. Inspection of genome browser tracks, harboring hundreds of active and inactive PINTS regions, confirm excellent concordance between H2BNTac with active PINTS regions (**Extended Data Fig. 17**). Collectively, these analyses show that the validation rate of H2BNTac^+^ candidate enhancers is comparable or higher than enhancers defined by other features^13,14,37-40.^

### Active distal enhancers lacking H3K27ac and/or H2BNTac are rare

Having confirmed a high specificity and validation rate of H2BNTac^+^ candidate enhancers, we aimed to get an estimate of H3K27ac^−^ non-canonical enhancers ^7-10,41^. We analyzed MPRA-defined enhancers in mESC^36^. We assumed that (1) enhancer activity in MPRA is not biased by the native enhancer chromatin modifications. (2) ATAC^+^MPRA^+^ regions represent a set of candidate enhancers that are likely active in vivo. (3) ATAC^+^MPRA^+^H3K27ac^−^ or ATAC^+^MPRA^+^H2BNTac^−^ regions the fraction of endogenously active non-canonical enhancers. Strikingly, 88% (9,249 out of 10,531) of distal ATAC^+^MPRA^+^ regions overlapped with H2BK20ac and/or H3K27ac. In the top25% highly active ATAC^+^MPRA^+^ enhancers, >93% overlapped with H2BK20ac and/or H3K27ac regions (**Fig. 6a**). The ATAC^+^MPRA^+^ regions that did not overlap with H3K27ac and/or H2BNTac had low chromatin accessibility (**Fig. 6b**), indicating that these enhancers are closed in the native context. This indicates that a vast majority of endogenously accessible and active enhancers are marked with H3K27ac and/or H2BNTac.

**Fig. 6.**
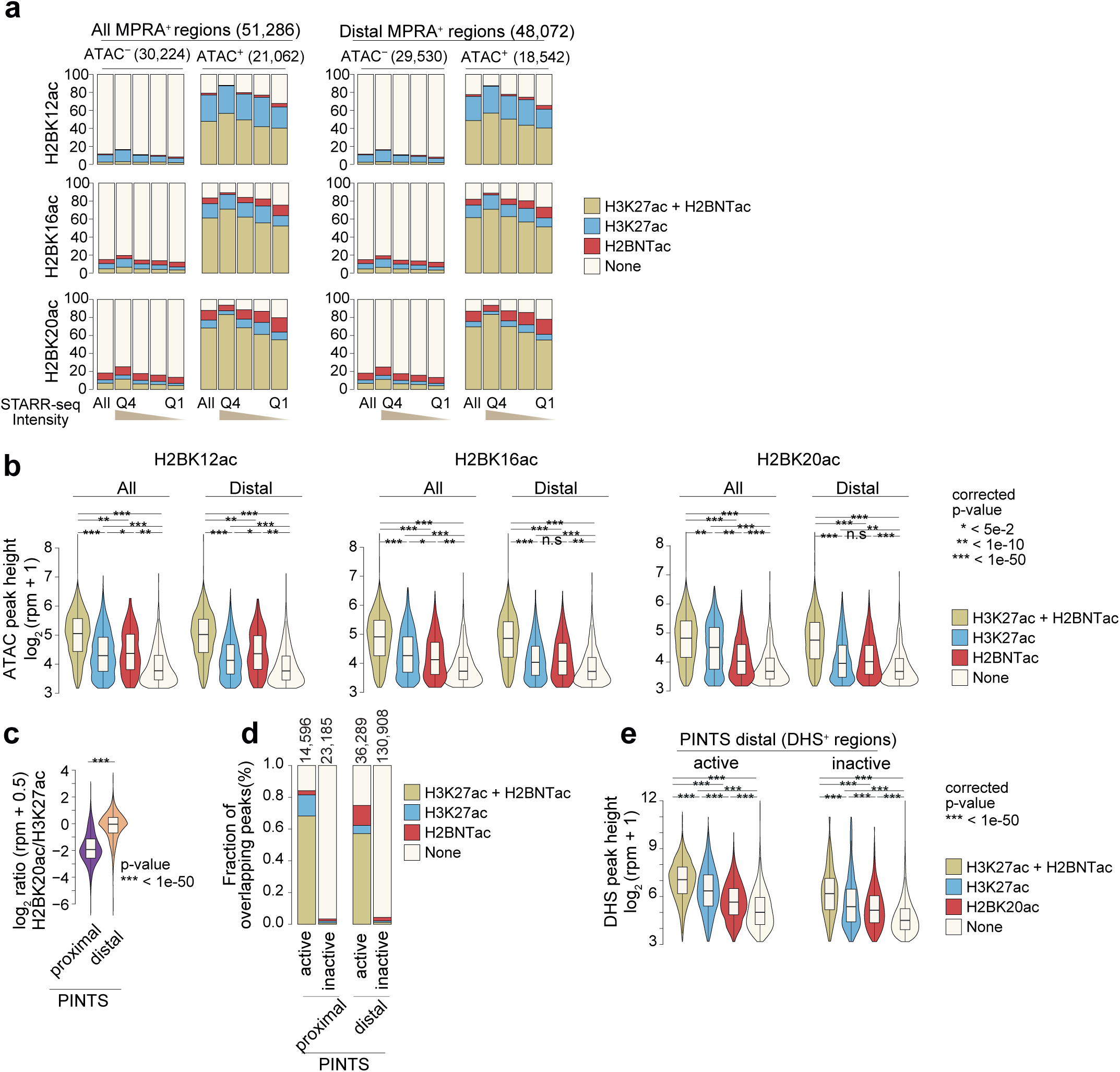
Most active enhancer regions are marked by H2BNTac and H3K27ac. **a,** A majority of ATAC+MPRA+ regions are marked by H2BNTac and/or H3K27ac. MPRA regions were grouped into ATAC^+^ and ATAC^−^. Within each group, MPRA^+^ regions were sub-grouped into quartiles based on the STARR-seq signal. Shown is the overlap of H2BNTac and/or H3K27ac in the indicated groups of MPRA regions. All: all MPRA^+^ regions; Distal MPRA^+^ regions: MPRA^+^ regions mapping >±1kb away from annotated TSS regions. **b,** H2BNTac and/or H3K27ac non-overlapping MPRA^+^ regions have low DNA accessibility. MPRA^+^ATAC^+^ regions were grouped based on the overlap of the indicated histone acetylation marks, and within each group, ATAC peak height is analyzed MPRA+-PRA^+^ATAC^+^ regions overlapping with the histone marks have much higher accessibility than regions lacking these marks. **c,** The relative H2BK20ac abundance of H2BK20ac is higher in distal PINTS regions. Shown is the ratio of H2BK20ac/H3K27ac ChIP-seq signal in proximal and distal PINTS regions in K562. **d,** A majority of PINTS active regions are marked by H2BNTac and/or H3K27ac. PINTS regions were grouped into proximal and distal, and further sub-grouped into active and inactive (see Methods-Shown is the fraction of PINTS regions overlapping with H2BK20ac and/or H3K27ac in K562. **e,** Distal PINTS active regions lacking H2BK20ac and/or H3K27ac have low DNA accessibility. Distal PINTS active regions were grouped based on the occurrence-of the indicated histone marks. Within each category, DHS peak height is determined. Distal PINTS active regions marked with H2BK20ac and/or H3K27ac have much higher accessibility than regions lacking these marks. Statistical significance determined by Mann–Whitney U test.

To independently verify this conclusion, we used PINTS-defined candidate active enhancers as a reference. We assumed that distal active PINTS regions marked with H3K27ac and/or H2BK20ac represent canonical enhancers, and regions lacking these marks indicate non-canonical active enhancers. The ratio of H2BK20ac/H3K27ac was significantly higher at distal PINTS regions than at proximal PINTS regions (**Fig. 6c**). 75% of distal active PINTS regions were marked by H3K27ac and/or H2BK20ac, whereas just 4% of distal inactive PINTS regions were marked by H3K27ac and/or H2BK20ac (**Fig. 6d**). Notably, non-overlapping active PINTS regions had low accessibility (**Fig. 6e**), indicating that most natively accessible enhancers are marked by H3K27ac and/or H2BK20ac. Together, these analyses suggest that H3K27ac and/or H2BNTac mark most of the endogenously active enhancers. Non-canonical active enhancers are either rare or occur in regions that lack or have very low DNA accessibility.

### H2BNTac correlates with enhancer-dependent gene regulation

Several promoter-associated features positively correlate with gene expression level^42^. We investigated the relationship between H2BNTac abundance, gene expression level, and cell-type specificity in mESC. Genes were rank-ordered based on MED1, H3K27ac, H3K9ac, and H2BNTac ChIP signals in promoter regions, and gene expression levels and cell-type-specificity were determined within each rank category. H3K27ac, H3K9ac, and MED1 abundance positively corresponded with gene expression, but H2BNTac showed no clear relationship (**Extended Data Fig. 18a**). Instead, unlike H3K27ac, H3K9ac, and MED1, promoter H2BNTac abundance was positively associated with cell-type specificity; genes with a higher H2BNTac signal were expressed in fewer cell types (**Extended Data Fig. 18b**). An A-485-induced decrease in promoter H3K27ac signal correlated with gene downregulation, but H2BNTac showed poor correlation (**Extended Data Fig. 19**), likely because H2BNTac is decreased globally whereas only a subset of genes is downregulated.

It would be most useful if the native abundance of a chromatin mark could predict a genés enhancer dependence. We noted that, in promoter regions of A-485-downregulated genes ^21^, the H2BNTac/H3K27ac ratio was significantly higher than in unregulated genes (**Fig. 7a**), indicating an association of promoter H2BNTac signal with CBP/p300-dependent gene regulation. To test this, genes were ranked based on MED1, H3K27ac, H3K9ac, and H2BNTac ChIP signal in promoters, and the fraction of A-485 downregulated genes was determined. Globally, ∼13-14% of genes were downregulated (≥2-fold) after 1h of CBP/p300 inhibition (**Fig. 7b**). Ranking of genes by MED1 and H3K9ac promoter signal showed no relationship with A-485-induced gene regulation. H3K27ac was modestly associated with enhancer-dependent gene regulation; among the top 100 genes, 35% of genes were downregulated. But, among the top 1,000 H3K27ac^+^promoters, only 20% of genes were downregulated, which is only marginally higher than the average (13%).

**Fig. 7.**
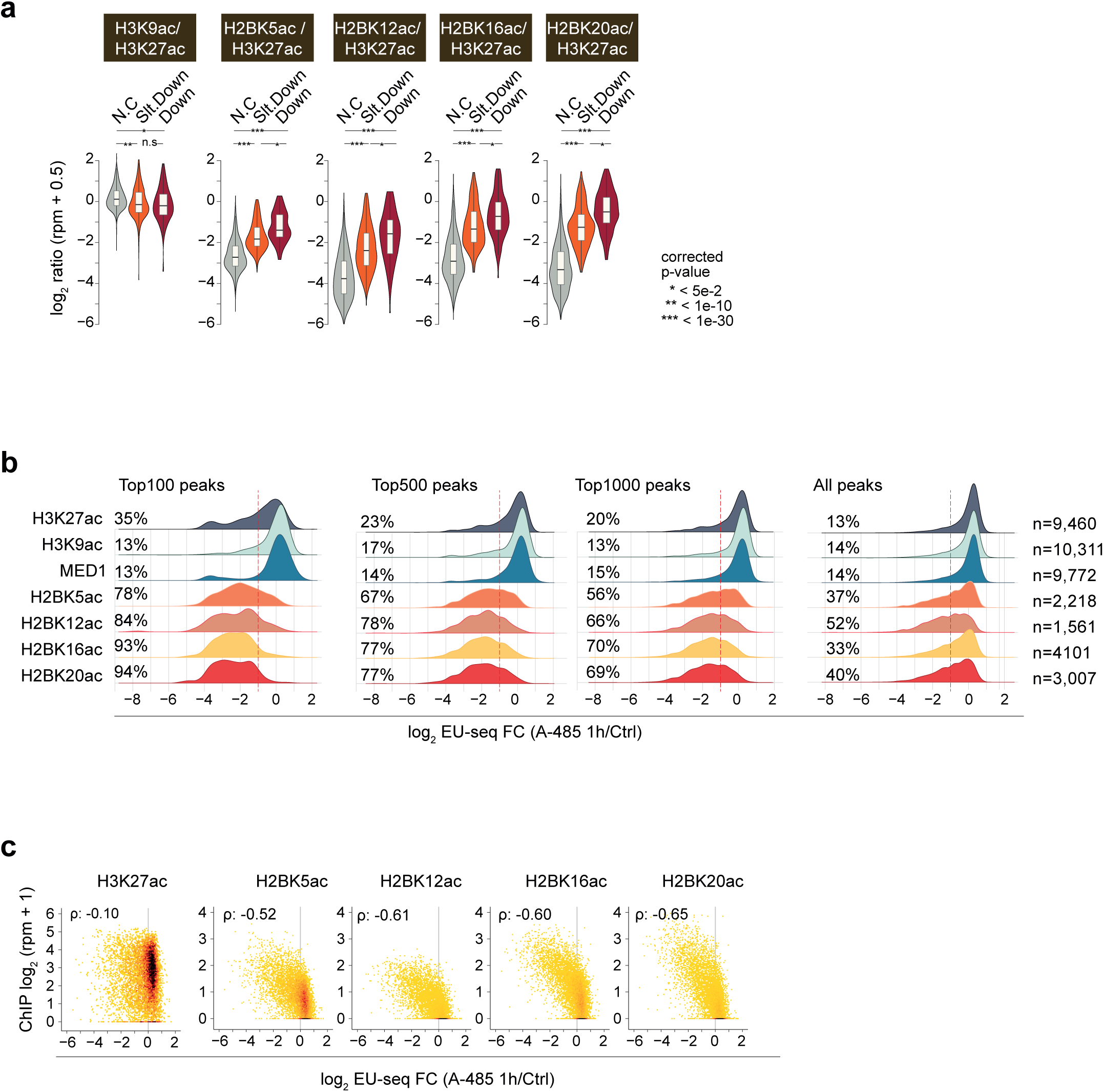
H2BNTac marks cell-type-specific promoters and predicts enhancer target genes in mESC. **a,** Shown is the ratio of H2BNTac sites and H3K27ac in the promoters of the indicated CBP/p300-regulated gene categories. mESC genes were grouped into A-485 down-regulated (Down), slightly downregulated (Slt. Down), or not changed (N.C.) categories, as defined ny Narita et al.^21^. The ratio of indicated histone marks was determined by normalized ChIP-seq counts in promoters (within ±1 kb of TSS) of the respective gene categories. Statistical significance determined by Mann–Whitney U test. **b,** A high H2BNTac promoter intensity predicts enhancer target genes. For the indicated chromatin marks, mESC genes are ranked based on the ChIP-seq signal in promoter regions (within ±1 kb of TSS). Shown are the composite nascent RNA transcription profiles for the top 100, 500, and 1,000 genes. Within each group, the fraction of A-485 downregulated genes is indicated. Change in nascent transcription was determined by EU-seq in mESC treated without or with A-485 (1h)^21^. The dotted lines indicate a ≥2-fold downregulation of nascent transcription after the A-485 treatment. **c,** Correlation between the ChIP-seq signal of indicated chromatin mark in promoter regions, and A-485 induced nascent transcription changes in mESC. (ρ: Spearman’s rho).

H2BNTac promoter intensity showed a stronger association with A-485-induced gene downregulation. In the top 100 highest H2BNTac marked promoters, 78-94% of genes were downregulated, in the top 500 genes, 67-78% were downregulated, and in the top 1,000, 56-70% were downregulated (**Fig. 7b**). Indeed, the overall distribution of H2BNTac marked genes is shifted towards downregulation and many H2BNTac positive genes were downregulated but fell below the 2-fold cut-off. This trend was confirmed in global analyses; A-485-induced gene downregulation correlated more strongly with H2BNTac than H3K27ac promoter intensity (Spearman’s ρ = −0.52 - −0.65 versus −0.10) (**Fig. 7c**).

### Promoter H2BNTac signal predicts enhancer target genes

Individually, chromatin features poorly predict enhancer targets. The activity-by-contact (ABC) model improves prediction accuracy by combining H3K27ac, DHS, and Hi-C contact frequency^43^. We compared the predictive performance of the H3K27ac and H2BNTac, with or without using the ABC model. Our positive set included genes that are highly downregulated (>2-fold) after CBP/p300 inhibition, and the negative set included genes that remained unaffected (<1.2-fold regulation) by CBP/p300 inhibition^21^. For each enhancer and gene, the ABC score is calculated pairwise. One gene can receive input from multiple enhancers. In this gene-centric approach, summed nominal ABC scores should reflect enhancer-dependent gene regulation.

Consistent with the original report^43^, the ABC model performed much better than simple H3K27ac promoter intensity (AUPRC:0.58, AUROC:0.76 versus AUPRC:0.31, AUROC:0.47) (**Fig. 8a-b**). The prediction accuracy of ABC model was greatly increased when we substituted the H3K27ac ChIP signal with H2BNTac (AUPRC: 0.62-0.64, AUROC: 0.78-0.79). In the original ABC model, the 3D contact information was obtained using Hi-C data. Because Micro-C provides better resolution than Hi-C^44,45^, we determined ABC scores using Micro-C and H2BNTac. Substituting Hi-C with Micro-C only marginally improved the model’s predictive power (**Fig. 8a-b**). 3D enhancer-promoter contact data are not available for many cell lines. Because H2BNTac promoter intensity correlated with enhancer-dependent gene regulation (**Fig. 7**), we tested the predictive value of H2BNTac intensity. Strikingly, H2BNTac promoter intensity alone, without involving DHS or 3D genome contact data, outperformed the ABC model and provided the highest prediction accuracy (AUPRC: 0.83-0.84, AUROC: 0.92) (**Fig. 8a-b**). Of note, despite limited ChIP-seq coverage, the performance of H2BK12ac was only marginally lower than H2BK16ac and H2BK20ac, indicating that sampling the ∼10,000 most abundant H2BNTac peaks can achieve a good prediction accuracy in mESC. Because of the lack of specificity of H3K27ac, the ABC model excludes the ChIP signal in promoter regions (+/-500bp from TSS). Our findings show that, if the ABC score is calculated using H2BNTac, excluding promoter intensity is disadvantageous for predicting enhancer targets. This implies that H2BNTac promoter intensity better reflects enhancer contribution to gene activation than H3K27ac and that the H2BNTac intensity can be used to estimate an enhancer activity score (A score). The ‘A score’ offers a simpler and more accurate tool for predicting strongly regulated enhancer targets and their dependencies.

**Fig. 8.**
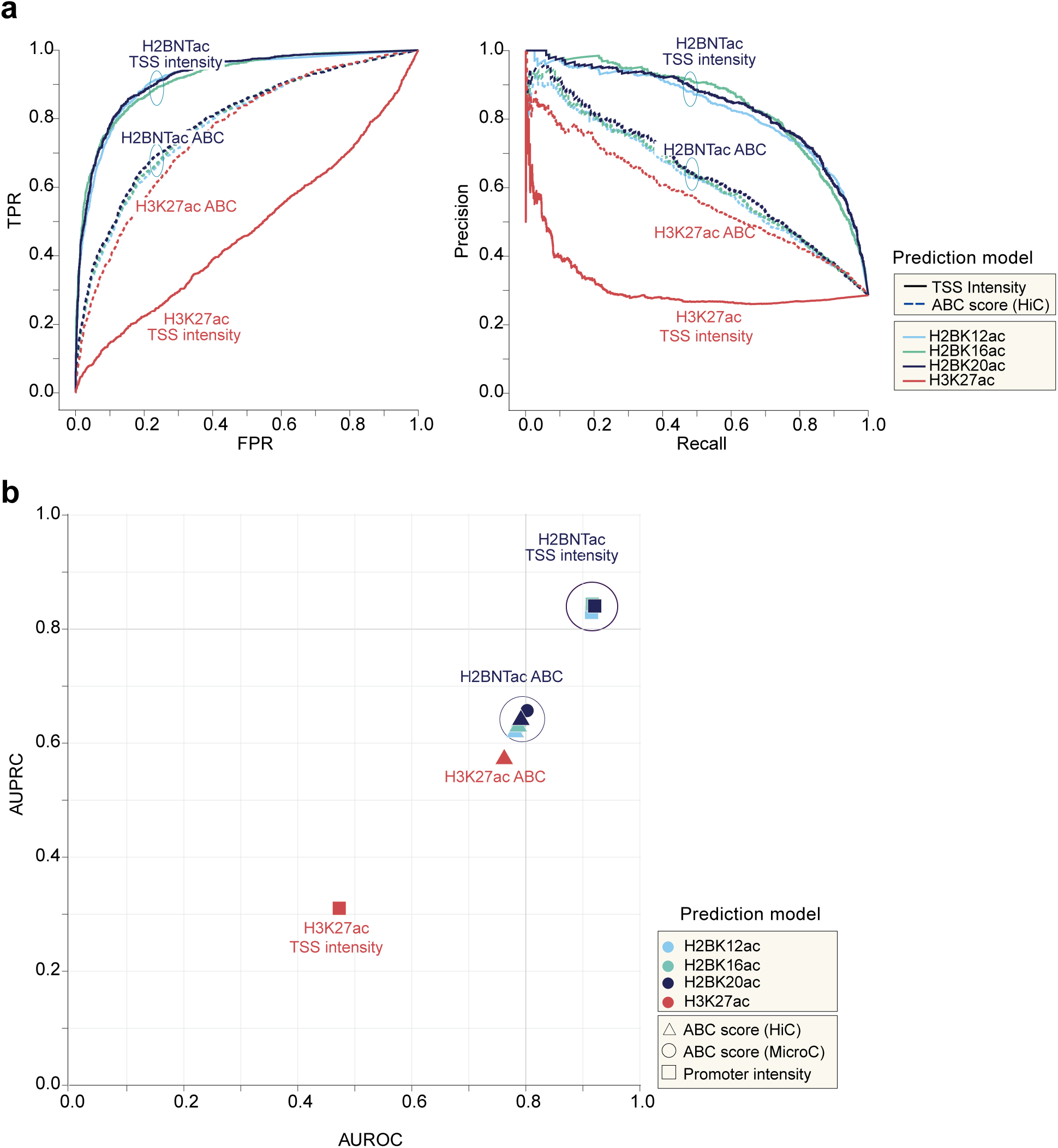
H2BNTac promoter intensity predicts enhancer-regulated genes in mESC. **a,** Receptor operative curve (ROC) showing the performance of the indicated histone marks in discriminating CBP/p300-dependent and -independent genes. CBP/p300-dependent (positive list) and independent genes (negative list) were defined by nascent transcription analyses after acute CBP/p300 inhibition^21^. Genes downregulated (≥2-fold) after A-485 treatment (average fold change of 30min, 1h and 2h treatment, see Methods) are considered enhancer-dependent, and unaffected genes (<1.2-fold change) are considered enhancer-independent. Performance was evaluated using cumulative nominal ABC scores or ChIP-seq enrichment of H3K27ac and H2BNTac marks in promoter regions (within ±1kb from TSS). **b,** H2BNTac promoter intensity outperforms H3K27ac and the ABC model in predicting enhancer target genes. AUPRC and AUROC were determined as indicated above in Fig. 8a. Shown are AUPRC and AUROC for the indicated histone marks. For H3K27ac, the ABC score is calculated by Fulco et al. using Hi-C contact frequency^43^. For the indicated H2BNTac sites, the ABC score was separately calculated using Hi-C or Micro-C contact frequency.

## Discussion

Through rational follow-up of the CBP/p300-regulated acetylome^18^, we rigorously establish H2BNTac as a distinctive and valuable signature of enhancers. Since H3K27ac was shown to mark enhancers^11,26^, the number of H3K27ac genome-wide mapping studies has increased linearly whereas H2BNTac has remained largely unstudied (**Extended Data Fig. 20**). This unambiguously shows that, until now, H2BNTac is not considered as a useful chromatin mark for defining CREs. Our findings suggest that this perception needs to be revised.

For the following reasons, we believe that the H2BNTac signature is robust and unlikely to arise from a lack of antibody specificity. (1) We used six different rabbit monoclonal antibodies, raised against 5 different H2BNTac sites, to establish the H2BNTac signature. (2) Because histone termini are unstructured, antibody specificity is primarily dictated by linear sequences, not 3D conformation. Cross-reactivity can arise if amino acids flanking modified residues are identical. (3) Within the H2BNT, amino acids flanking lysine are highly dissimilar, making it unlikely that all monoclonal antibodies cross-react within H2BNT. (4) Guided by the sequence similarity, and the unique H2BNTac profile, we found one of the H2BK5ac antibodies as an outlier, explained its cross-reactivity with H3K27ac, and excluded it from our analyses. (5) Transcription inhibition preferentially increased H2BNTac in actively transcribed genes, consistent with the transcription-induced exchange of H2A-H2B^32,33^. (6) CBP/p300-dependent H2BNTac regulation in our ChIP-seq is consistent with mass spectrometry-based analyses of H2BNTac sites^18^. The only other histone sites that show similarly strong, global downregulation after CBP/p300 inhibition are a subset of sites present in histone H3 N-terminus, whose genome occupancy profiles are dissimilar from H2BNTac. (7) CREs marked with H2BNTac bear all the canonical features of active enhancers, including DHS, H3K4me1, H3K27ac, and MED1 binding. (8) H2BNTac marked regions show a high validation rate in orthogonal enhancer activity assays.

To our knowledge, this work presents the first coherent, genome-scale map of H2BNTac site occupancy and locus-specific regulation. Our analyses confirm the enhancer specificity of H2BK20ac^9^, but we found no concrete evidence of a distinct class of strong H2BK20ac^+^ enhancers that lack H3K27ac^9^. In our analyses, incomplete overlap most likely reflects the technical difficulty in reproducibly detecting low abundant peaks. Our analyses conclusively show that H2BK20ac is not an outlier among H2BNTac sites, and unlike acetylation sites in the N-termini of histones H3 and H4, all analyzed H2BNTac sites almost indistinguishably mark the same genomic regions. Furthermore, for the first time, our analyses offer mechanistic reasons for the distinct H2BNTac and H3K27ac signatures and identify H2BNTac as the most specific diagnostic marker of CBP/p300 activity. We reveal a yet underappreciated role of transcription-coupled nucleosome exchange in shaping H2BNTac occupancy^32,33^. This implies that, unlike modifications on H3 and H4, modifications on H2A and H2B in actively transcribed regions may, in general, be short-lived. The differential exchange of H2A-H2B and H3-H4 in TSS upstream and downstream nucleosomes through transcription-coupled nucleosome exchange ^33^ may also explain the asymmetric abundance of H3-H4-associated marks, such as H3K4me3, H3K9ac, and H3K27ac, in the +1 and -1 nucleosomes.

Beyond providing a unified picture of H2BNTac, we used the distinctive H2BNTac signature to address two key questions that are frequently discussed in contemporary review articles^12,17,41,46^. (1) How reliable are histone acetylation marks for predicting candidate enhancers? (2) What fraction of mammalian candidate enhancers remain undetected by histone acetylation marks?

We tested the reliability of H3K27ac and H2BNTac, using candidate enhancers defined by MPRA and eRNA transcription. 79% of all H2BNTac^+^ regions show evidence of RNA transcription, and among the top 25% regions, virtually all express RNA (**Figure 5b**). This shows that almost all highly acetylated regions in mammalian cells are actively transcribed. The differences in validation rate are associated with histone mark abundance and the depth of analyses. The validation rate among the top 25% of highly acetylated regions is higher than the bottom 25% regions. Similarly, if the depth of ChIP-seq analyses is limited, the validation rate appears higher. For example, a greater fraction of H2BK12ac^+^ regions are validated than H2BK20ac^+^ regions (**Figure 5a**), likely because the number of identified H2BK12ac peaks is lower than H2BK20ac. Similarly, with a ∼70% validation rate, eRNA was suggested to be a better surrogate of active enhancers than H3K27ac^13^, but with increased depth, the validation rate of eRNA-defined enhancers decreases from ∼70% to ∼30% ^14^, which is no higher than the validation rate of candidate enhancers defined by histone marks. The same logic (i.e. differences in the depth of analyses and sensitivity of assays used) can rationalize the variable validation rate in other prior studies^13,14,37-40^. Overall, we conclude that histone acetylation marks are reasonably sensitive and accurate in sampling a large fraction of active enhancers. An advantage of MPRA and eRNA-centric approaches is that they offer higher resolution and can more precisely map enhancer position than histone mark peaks, which tend to be broader. Histone modifications have the practical advantage that DNA is more stable than RNA, and ChIP-seq is relatively cheaper and less laborious.

Mammalian cells are suggested to harbor different classes of distal enhancers ^9,10,22,23,41^. If non-canonical enhancers were widespread, we would expect to find many ATAC^+^MPRA^+^ regions without H3K27ac^−^. We used thousands of ATAC^+^MPRA^+^ and eRNA-defined enhancers to get an approximation of non-canonical enhancers. 79-88% of ATAC^+^MPRA^+^ defined enhancers, and 79% of active PINTS^+^ regions, overlap with regions harboring H3K27ac and/or H2BNTac. The remaining non-overlapping regions have low chromatin accessibility, indicating that these enhancers are closed in the native context (**Fig. 6**). Some closed enhancers can bind pioneering TFs, such as p53, and can activate transcription in MPRA^36,47^, but they too acquire H3K27ac upon their activation in the native context^36^. We do not rule out the existence of non-canonical active enhancers without H3K27ac^−^ and/or H2BNTac^−^, but in our analyses, we did not find strong evidence for a large number of strong, endogenously active enhancers that lacked H3K27ac and/or H2BNTac. This realization is important for estimating the extent to which canonical and non-canonical enhancers likely contribute to global gene regulation.

One of the salient findings of our work is that H2BNTac is a useful marker for predicting enhancer target genes. This demonstrates that H2BNTac is not just ‘yet another enhancer marker’ but it offers genuinely complementary advantages to the existing marks. A notable advantage of H2BNTac is that it does not require 3D contact information for predicting enhancer targets, which makes it attractive for using in diverse cell types and tissues. While the dispensability of 3D contact information for predicting enhancer targets seems surprising at first glance, this observation is consistent with the observation that 3D genomic contacts remain largely unchanged during differentiation, even though gene expression is highly rewired^48,49^.

This, in no way, should be construed as E-P contact frequency being irrelevant; rather, it highlights the difficulty in faithfully capturing E-P contacts that may be weak and highly dynamic. We must also acknowledge that the overall correlation (p=-0.52--0.65) between promoter H2BNTac and gene regulation is not perfect and there is room for future improvements. In the accompanying work, we show that the functional impact of enhancers is affected by the E-P distance, as well as the autonomous activation strength of promoters (see the accompanying manuscript, Narita et al.). Thus, including the distance calibrated distal H2BNTac signal, and factoring in autonomous promoter strength could present future avenues for further improving the predictive potential.

While our work rigorously establishes the specificity of H2BNTac, the biological relevance of H2BNTac remains to be investigated. This task is tedious because H2B is encoded by multiple genes, and multiple lysines in the N-terminus are modified by multiple modifications, possibly serving distinct functions. Hypothetically, a string of dynamically regulated H2BNTac has the potential to exert functional influence by affecting histone-DNA interaction^50^, influencing the phase-separation property of chromatin^51^, or serving as a ligand for bromodomain proteins, which preferentially interact with multiply acetylated peptides^52^. At this point, these are mere ideas, and further work is required to investigate them. To avoid confusion, here we present no claim, explicit or implied, that H2BNTac is functionally more important than other histone acetylation marks. Because H3K27ac is dispensable for gene activation^53-57^, we suggest that the CBP/p300 function in transcription activation should be considered beyond H3K27ac, and H2BNTac merits consideration.

Collectively, our findings provide a unified view of H2BNTac genomic occupancy and its locus-specific regulation. Identification of H2BNTac as a distinctive active enhancer signature will facilitate fine-grained mapping of CREs, better prediction of enhancer strength and function; and thereby, contribute to improved understanding of gene regulation. In the accompanying manuscript, we exemplify how unique H2BNTac attributes can enable an improved understanding of native enhancer-promoter compatibility, and expose hitherto obscure relationships between enhancer strength, distance, and gene regulation.

## Supplemental note 1

For more than 2 decades, acetylation of multiple H2BNT sites has been detected across diverse eukaryotes^58,59^, even though the H2BNT sequence shows little sequence conservation between budding yeast, *Drosophila*, and human (**Extended Data Fig. 20b-c**). The current picture of H2BNTac genomic occupancy and regulation is confusing. In three different cell lines, H2BNTac sites showed varying correlations with each other and with other chromatin marks^9,22,23^. H2BNTac sites formed a 17-histone modification signature that marked virtually all active promoters^22^. Similar to H3K27ac and H3K4me3, promoter abundance of H2BK5ac was strongly correlated with global gene expression^42^. One study noted preferential enrichment of H2BK20ac at active enhancers and cell-type-specific promoters; but most notably, this study identified an entirely new class of enhancers that were exclusively marked with H2BK20ac and lacked the canonical enhancer mark H3K27ac^9^. Furthermore, H2BK20ac occupancy correlated with H2BK120ac, but not with other H2BNTac sites^9,23^, suggesting that it is an outlier among H2BNTac marks. More recently, H2BK20ac was shown to be critical for recruiting histone macroH2A1, which then recruits PARP1 and CBP to promote acetylation of H2BK12 and H2BK120^60^. Contrasting these findings, proteomic analyses showed that virtually all H2BNTac sites were similarly regulated by CBP/p300^18^, that the abundance of H2BK20ac is orders of magnitude higher than H2BK120ac^20^, and that, unlike H2BK20ac, H2BK120ac is neither impacted by CBP/p300 nor by any of broadly used deacetylase inhibition^18,19^.

Together, the prior literature portrayed a complex picture of H2BNTac occupancy and left many mechanistic questions unresolved. For example, why is H2BK20ac an outlier among H2BNTac sites, and why do other H2BNTac sites lack enhancer specificity? How does the same acetyltransferase, CBP/p300, differentially acetylate H3K27 and H2BK20 in a locus-specific manner? Is H2BK20ac catalyzed by acetyltransferase other than CBP/p300 at the new class of H2BK20ac^+^H3K27ac^─^ enhancers? Why does H2BK20ac genome occupancy resembles that of H2BK120ac, are these marks differentially regulated by acetyltransferase and deacetylases? The sequential H2BK20 and H2BK120 acetylation model^60^ could explain why these marks co-occur, but it also suggests that H2BK20ac is already present before the recruitment macroH2A1-PARP1-CBP/p300, and H2BK12ac occur after that. What acetylates the initial H2BK20 before the recruitment of macroH2A1, and if PARP1-CBP acetylates H2BK12 and H2BK120 in the same regions^60^? If so, why do these two marks show dissimilar genomic occupancy^22,23^? How do H2BK20ac and other H2BNTac sites compare with other chromatin features in the prediction of enhancer target genes?

The lack of mechanistic understanding emerges as the primary reason for these discrepancies, which, combined with poor H2BNT sequence conservation, has hampered the adoption of H2BNTac as a reliable enhancer marker. Our prior mass spectrometry analyses clarified the site-specificity and dynamics of CBP/p300 targets^18^. We harnessed this knowledge to uncover and explain a unifying picture H2BNTac sites. This illustrates the usefulness of integrating proteomic and genomic analyses in obtaining an improved mechanistic understanding of chromatin marks.

## Acknowledgments

We thank the members of the Choudhary lab for their helpful discussions. We thank Elina Maskey for her excellent technical assistance. The Novo Nordisk Foundation Center for Protein Research is financially supported by the Novo Nordisk Foundation (NNF14CC0001). C.C. was supported by the Novo Nordisk Foundation (NNF14OC0008541). This work was funded by the European Commission FP7 grant (SyBoSS FP7-242129). Y.H. was supported by a Grant-in-Aid for JSPS Overseas Postdoctoral Fellows. S.K. was supported by a Lundbeck Foundation Fellowship (R347-2020-2170). We thank the SyBoSS partners Konstantinos Anastassiadis, A. Francis Stewart, Austin Smith, and Bill Skarnes for sharing E14TG2a Oct4-IRES-Puro mouse ESCs. We thank Abcam, Cell Signaling Technology, Thermo Fisher Scientific, Active Motif, and RevMAb Biosciences, for providing some of the antibody samples for testing. We thank the CPR Imaging Platform, the CPR Big Data Management Platform, and the CPR and DanStem Genomics Platform for their assistance. We thank the ENCODE Consortium and the ENCODE production laboratory for generating and sharing the data.

## Author contributions

C.C. and T.N. conceived the project. T.N., Y.H., S.K., T.L. J.W. designed research, performed experiments, analyzed data, and interpreted the results. C.C. supervised the project. T.N. and C.C. wrote the manuscript with input from all co-authors.

## Conflict of interest

The authors declare no conflict of interest.

**Supplementary Table 1** A list of H2BNTac antibodies tested and used for ChIP. Before using for ChIP-seq, performance of all antibodies was tested using ChIP-qPCR. From chromatin immunoprecipitated samples, fold enrichment of acetylation-positive (*Nanog* enhancer) and acetylation-negative chromatin region (*Hoxa13*) regions was determined using ChIP-qPCR. Antibodies providing >5-fold enrichment were used for ChIP-seq analyses.

**Supplementary Table 2** Sequences of unmodified and acetylated histone peptides that were used for analyzing antibody specificity by quantitative image-based cytometry.

## Materials and methods

### Cell culture

E14TG2a Oct4-IRES-Puro mouse ESCs^64^ were cultured in a custom-made (C.C.Pro GmbH, Oberdorla) N2B27 medium consisting of a 1:1 mix of DMEM/F12 and Neurobasal medium, but lacking Arginine and Lysine. Before use, the media was supplemented with 1000 units/ml leukemia inhibitory factor (LIF) (Merck Millipore), 1µM PD0325901, 3µM CT-99021 (custom-made by ABCR GmbH, Karlsruhe), 100µM 2-mercaptoethanol, 150µM sodium pyruvate, 0.5x B27 supplement (Thermo Fisher Scientific), 0.5x N2 supplement (made in house, or from Thermo Fisher Scientific), and where relevant with L-lysine and L-arginine. Where indicated, the cells were treated with A-485 (10µM). K562 and RPE1 cells were cultured in DMEM media supplemented with 10% FBS, and 1% penicillin/streptomycin. All cells were cultured at 5% CO2, 37 oC.

### EU-seq differential gene expression analysis (DGE)

Previously published (GSE146328)^21^ EU-seq data of ESC treated with DMSO or A-485 (10µM, 30, 60, and 120 min) were re-processed. Adaptor and low-quality sequence trimming was performed as described in the RNA-seq section. Read sequences were aligned to mm10 (mouse) using bwa aln with default parameters (BWA version 0.7.10^65^. Multi-mapped reads and reads with more than three mismatches were removed using samtools (version 1.4)^66^. Reads mapped to rRNA and tRNA region obtained from UCSC genome browser were removed using Bedtools. For 30, 60, and 120 min timepoint data, based on the Pol2 elongation rates, maximum 30, 90, and 240 Kb gene body regions from TSS were used for differential gene expression (DGE) analysis. The number of reads mapped to defined regions was counted using HTseq. Log2 fold-change (FC) and p-values were calculated in individual time points using DEseq2^67^ with the default scaling method (median of relative abundance). Expressed genes were determined as described in the reference genome annotation section and further filtered the low expressed genes if the mean EU-seq read count of control and A-485 treatment condition is less than 20.

### ChIP

The cells were cross-linked with 1% formaldehyde for 10 min. After 5 min neutralization with 0.2 M glycine, the cells were collected, re-suspended in SDS lysis buffer, composed of 10 mM Tris-HCl, pH 8.0, 150 mM NaCl, 1% SDS, 1 mM EDTA, pH 8.0, and cOmplete™ EDTA-free Protease Inhibitor Cocktail (Sigma-Aldrich), and fragmented by a Bioruptor Pico (10 cycles, 30 sec on/30 sec off; Diagenode). The sonicated solution was diluted (1:5 ratio) with ChIP dilution buffer (20 mM Tris-HCl, pH 8.0, 150 mM NaCl, 1 mM EDTA, 1% Triton X-100) and then used for ChIP. 2-5 μg of antibody was bound to pre-blocked (with 0.5% BSA) magnetic Dynabeads M-280 sheep anti-Mouse IgG or sheep anti-Rabbit IgG (Thermo Fisher Scientific) and applied to the diluted, sonicated solution for ChIP. Chromatin was added to the antibody-bead complex and incubated rotating overnight at 4°C. The beads were washed with ChIP dilution buffer, wash buffer high salt (20mM Tris-HCl, pH8.0, 500mM NaCl, 2mM EDTA, 1% Triton-X, 0.1% SDS), wash buffer low salt (10mM Tris-HCl, pH8.0, 1mM EDTA, 250mM LiCl, 0.5% Na-Deoxycholate, 0.5% NP-40) and TE buffer. The bound materials were eluted with elution buffer (50mM Tris-HCl, pH8.0, 10mM EDTA, 1% SDS) overnight at 65°C and treated with RNaseA for 30 min at 37°C and further incubation with Proteinase K for 1 hour at 55°C. Then DNA was purified with phenol-chloroform extraction.

### ChIP-qPCR

To validate acetyl-histone antibodies for ChIP-seq, we performed ChIP-qPCR to measure the fold-enrichment of modified chromatin. Approximately equal amounts of chromatin was mixed with acetyl-histone antibodies listed in Supplementary Table1, and then the relative enrichment of positive (Nanog enhancer) and negative (Hoxa13) genomic regions was quantified by ChIP-qPCR. Antibodies giving fold-enrichment (Nanog/Hoxa13) >5 were used for ChIP-seq analysis. Nanog enhancer region (Fw: CTTGGGAGAGAGGGAAAGAAACA, Rv: AGTCAGGACCTCACTATGTCAGA) and Hoxa13 intron (Fw: CACCAAATTGTCCCTGATGGTTC, Rv: ACACTTCTTTCTGTAGAGCTCGG) are used as primers.

### ChIP-seq library preparation

ChIP-Seq library was prepared using DNA sonicated to an average size of 0.5 kb. ChIP samples were processed for library preparation using the NEBNext Ultra II DNA Library Prep Kit for Illumina (NEB) following the manufacturer’s instruction and sequenced on a Next-Seq 500 Sequencer (Illumina) as single-end 75b reads.

### Processing of ChIP-seq data

Mouse and Human genome annotation and reference genomes were downloaded from the GENCODE website (mouse; GRCm38 release 25, human; Grch37 release 29) ^68^. Transcript type "protein_coding" was chosen as representative genes. For genes with multiple isoforms, the longest isoform was used for the analyses. Reads were mapped to the reference genome by using bwa aln with default parameters (BWA version 0.7.10) ^65^. Multi-mapped reads, duplicated reads, or reads with more than 3 mismatches were removed by samtools ^66^. Reads mapped to the DAC Blacklisted Regions (https://www.encodeproject.org/annotations/ENCSR636HFF/) were also omitted from the downstream analysis. The following publicly available datasets were downloaded from the GSE repository and re-processed for consistency. ESC H3K27ac (GSE135562, GSE160890), K562 H3K9ac (GSE29611), K562 H3K4me3 (GSE163049) and K562 H3K4me1 (GSE29611, GSE31755).

Peak regions were called using LanceOtron with the default model (wide-and-deep_jan-2021) ^35^. Proximally occurring peaks (within 2 kb) are merged using Bedtools ^69^, and poorly enriched peaks of height < 8 reads per kilobase per million mapped reads (RPKM) were omitted. Peak height was calculated by bamCompare ^70^ with the following parameters (centerReads, minMappingQuality 10, 20bp bin, smooth length 400bp, extend reads 200bp, rpkm normalization and input subtracted). The region with maximum peak height within each peak loci was defined as a peak summit region. Where indicated, peak regions were classified into the following classes: “Promoter” (TSS ±1 kb), “Gene body” (Exon, intron, 5′ UTR, and 3′UTR, excluding “Promoter” regions), and “Intergenic” (outside promoter and gene body) by using ChIPseeker^71^. Mouse ESC super-enhancer region bed file was downloaded from dbSuper^63^, and converted from mm9 to mm10 using UCSC LiftOver function^72^. For the visualization of the ChIP signal, bigwig files were generated by bamCompare with the following parameters (centerReads, minMappingQuality 10, 20bp bin, smooth length 400bp, extend reads 200bp, rpkm normalization and input subtracted). For the visualization of ATAC and EU-seq signal, bigwig files were generated by bamCoverage with the following parameters (centerReads, minMappingQuality 10, 20bp bin, smooth length 400bp, extend reads 200bp, rpkm normalization). The Integrated genome viewer (IGV) is used for the visualization of gene tracks.

### ChIP-seq coverage analysis

In ChIP-seq coverage analysis, genomes were binned by a 2kb window and ChIP-seq read count was calculated using HTseq^73^. Read number was normalized by using rpm (reads per million mapped reads), and input subtracted values were used. For regions with negative values, the values were substituted with 0. Unless indicated otherwise, for comparing peak enrichment in the defined peak regions, a 2kb window of input-subtracted RPM value >1 was chosen, and a log2 ratio of RPM + 0.5 was used for calculating differential ChIP coverage ratios. For the analysis of promoter, promoter regions were defined as regions in ±1 kb around H3K4me3 marked TSS of actively transcribed genes (TPM ≥2), instead of genome-wide 2 kb bin. TPM values are calculated from EU-seq data (for mESC [GSE146328])^21^ or RNA-seq data (for K562 cell line).

### Peak overlap

In counting the overlap between ATAC peak, H3K27ac and H2BNTac peak regions, findOverlaps function (GenomicAlignments v1.8.4)^74^ with allowing maximum 2Kbp gaps was used. The peak number within overlapping regions are not identical between two peak sets because several peaks in one set can overlap with one peak in another set. The overlapping peak numbers shown in the figures are peak numbers of H2BNTac (when comparing with H3K27ac or ATAC peaks), or peak numbers of a set with more overlapping peaks (when comparing within H2BNTac). Wilcoxon rank-sum test was used for testing the significance of peak height difference between overlapping and non-overlapping peaks. In annotating peaks using OCT4 and NANOG bindings (Extended Data Fig S6), STARR-seq and ATAC peaks, PINTS region, and DHS regions, no gaps were allowed to be overlapped.

### Enrichment of ChromHMM states in peak regions

ESC 18 chromHMM states and K562 25 chromHMM states were downloaded from the encode portal^61^, and UCSC genome browser (http://hgdownload.cse.ucsc.edu/goldenpath/hg19/encodeDCC/wgEncodeAwgSegmentation) ^34,62^, respectively. We assigned chromatin states at each peak summit region as representative peak states. In the enrichment analysis, the expected peak number in each state was calculated by assigning an equal number of randomly sampled genomic regions, and the enrichment of chromatin states is defined by the ratio of the observed peak number to the expected peak number.

### Overlap of PINTS regions with peak regions

Annotations of proximal and distal PINTS elements for human cells were downloaded from the PINTS web portal (https://pints.yulab.org/summary_stats) and lifted over to hg19 using UCSC LiftOver function^72^. Using these data, we generated a reference dataset of PINTS regions identified in 110 cell lines and and tissues and merging proximally overlapping regions. We classified the PINTS regions into active and inactive in K562 cells using the following criteria. (1) Proximal active: proximal PINTS regions of K562 cells. (2) Proximal inactive: proximal PINTS regions that are active in other cell types but not in K562, and occurred >2kb away from any K562 proximal active PINTS regions. (3) Distal active: distal PINTS regions that are active in K562 cells. (4) Distal inactive: distal PINTS regions that are active in other cell types but not active in K562. The following inactive regions were excluded: (i) inactive regions that overlapped with any of the proximal PINTS regions in K562, (ii) inactive regions located within 5kb of active TSS in K562, and (iii) inactive regions located within 2kb from distal active PINTS regions in K562. Overlap between PINTS regions with ChIP-seq peak is calculated using distanceToNearest function (GenomicRanges v1.30.1)^74^.

### Discriminative power assessment of PINTS regions

For each set of positive and negative PINTS regions, enrichment of the ChIP signal was calculated from input-subtracted RPM values. We computed the average ChIP signal by varying the window (+/-250, 500, 1000bp from the PINTS region center). The discriminatory potential of different histone marks was analyzed by comparing ChIP signal enrichment at different PINTS regions. Receiver operating characteristics (ROC) and precision-recall curves (PRC) were generated under the different thresholds of peak enrichment or peak enrichment ratios. ROC, PRC, and Area under the curve (AUC) were computed using PRROC R package ^75^.

### Discriminative power assessment of enhancer-regulated genes

H3K27ac-based ABC score was downloaded from the data availability section of Fulco et.al.^43^. H2BKNTac-based ABC scores were calculated using the ABC model pipeline (https://github.com/broadinstitute/ABC-Enhancer-Gene-Prediction). As input data for the ABC model, ATAC seq data (GSE146328), Hi-C (GSE118911), or Micro-C (GSE130275) were used for calling candidate regions and quantifying enhancer-target accessibilities, respectively. Hi-C data is reprocessed to map to the mm10 genome by using juicer ^76^ with filtering of MAPQ >30. Processed Micro-C data was downloaded from GSE repository (GSE130275_mESC_WT_combined_2.6B.hic). To calculate the ABC model-based enhancer contribution, nominal scores (powerlaw.Score.Numerator) were summed up for each target gene. For the genes with no ABC scores or a score < 0.01, an ABC score of 0.0.1 was imputed. In calculating the enhancer contribution based on the histone mark intensities, The ChIP reads enrichment at the promoter regions (±1Kbp from TSS) was used. As a positive and negative set of enhancer-regulated genes, >2 fold down-regulated genes and < 20% changes upon A-485 treatment based on EU-seq were selected. For a fair comparison, genes included in the processed data by Fulco et.al were analyzed. ROC and PRC were generated by using different thresholds of peak enrichment or cumulative ABC score. ROC, PRC, and Area under curve (AUC) were computed using PRROC R package ^75^.

### Statistical analysis

Otherwise mentioned, p-values are calculated using Wilcoxon rank-sum test and corrected for multiple comparisons using Benjamini & Hochberg method (R package stats version 3.6.2).

### Analysis of publically-available histone acetylation ChIP-seq projects in GEO

The openly-accessible ChIP-seq data was retrieved from NCBI GEO using R package reutils using the query “ "peak"[All Fields] AND gse[Filter] AND Genome binding/occupancy profiling by high throughput sequencing[Filter]”, where "peak" was replaced with histone acetylation marks (i.e. H3K27ac, H2BK5ac, H2BK11ac, H2BK12ac, H2BK15ac, H2BK16ac or H2BK20ac). Data deposited from the year 2009 to 2021 were retrieved. After a manual check of the duplicates and availability of datasets, the number of projects belonging to each dataset is counted by the year when the data become public.

### Acetylation of histone peptides

Unmodified peptides corresponding to histones H2B and H3 N-termini were synthesized (Schafer-N) with C-terminal biotinylated lysine. The peptide was dissolved in PBS to a concentration of 2mg/mL. Fully acetylated forms of H2B and H3 N-termini peptides were generated by in-vitro chemical acetylation with Sulfo-NHS acetate. 200µg peptide was mixed with 40µL (20µg/µL) Sulfo-NHS acetate in acetonitrile and incubated for 30min at RT. Excess NHS acetate in the reaction was quenched by the addition of Tris-HCl pH 7.5 to a final concentration of 100mM. To check for completion of the reaction, 2µL of the product was acidified and desalted with C18 stage-tips, eluted, dried, and re-dissolved in water with 0.1% formic acid. Acetylation was confirmed by mass spectrometry (Orbitrap Exploris 480 mass spectrometer). Sequence information of histone peptides used for analyzing antibody specificity is provided in Supplementary Table 2.

### Immunostaining

hTERT RPE-1 (ATCC: CRL-4000) were seeded onto 12mm coverslips distributed in a 6cm dish and allowed to attach overnight. The day after, cells were fixed with 4% formaldehyde in PBS for 15min and washed once with PBS. Coverslips were placed into a 24-well plate. Cells were permeabilized with 0.5% Triton X-100 in PBS for 5min followed by blocking in antibody diluent (sterile filtered DMEM with 10% FBS and 0.02% sodium azide) for 30min. The primary antibody was diluted 1:1000 in the antibody diluent and split into three 200µL stocks. In one, no additive was included, in the second 0.2µL of 0.67mg/mL unmodified histone peptide was added and in the third 0.2µL of 0.67mg/mL acetylated histone peptide was added. Two sets of coverslips for each antibody were then stained for 2h with the antibody without/with peptide. The coverslips were washed thrice with PBS and stained with secondary anti-rabbit Alexa 488 antibody diluted (1:500) in the antibody diluent for 1h. The coverslips were washed once and stained with Hoechst 33342 (1µg/mL) in PBS for 10min followed by two PBS washes, dipping into the water, and mounting on slides using 5µL Fluoromount-G.

### Image-based cytometry

Images for image-based cytometry were acquired on an Olympus ScanR automated widefield screening microscope with 12-bit dynamics on a 16-bit Hamamatsu ORCA-Flash4.0 2048x2048 pixel camera with 6.5µm and pixel size using a UPLSAPO 20x objective with 0.75NA. Images were analyzed using Olympus ScanR Image Analysis software version 2.8. Nuclei segmentation was carried out with the integrated intensity-based object detection using the Hoechst signal and after background correction, the total and mean intensities of the two channels were calculated for each object. Further analysis was done on properly segmented cells gated for having a 2C-4C DNA content based on total and mean DAPI intensities. Data tables from ScanR Analysis were further processed and visualized in R using ggplot2. 3000 randomly sampled cells from each coverslip were included for each experiment and repetition. Example images from each coverslip with set contrast and brightness settings for each antibody are displayed with their corresponding quantifications.

### Generation of HDAC1/2-GFP-FKBP12^F36V^ cells

GFP-FKBP12^F36V^ tag was knocked-in at the C terminus of the endogenous *Hdac1* and *Hdac2* using the CRISPR/Cas9 system. Mouse ESCs were co-transfected with pX330 plasmid and a donor plasmid containing resistance selection gene (*Hdac1*: Puro, *Hdac2*: Neo), flanked by homology arms (∼500bp each side from the start of the stop codon of the target genes). The sequences of guide RNAs for targeting *Hdac1* and *Hdac2*. Puromycin or neomycin-resistant cell clones were screened by genomic PCR using KOD Xtreme hot-start DNA polymerase (Millipore).

### Confirmation of HDAC1/2-GFP-FKBP12^F36V^ depletion

Cells were seeded at 15,000 per well in Perkin Elmer Cell Carrier Ultra PhenoPlate 96-well, black, optically clear flat-bottom plates coated with ThermoFisher *Geltrex*™ LDEV-Free Reduced Growth Factor Basement Membrane Matrix. After seeding, cells were grown for two days. Cells were treated with either dΤAG13 (100nM) or DMSO as solvent control. Finally, cells were fixed using 4% PFA in 1x PBS and blocked with 5% BSA and 0.3% TritonX in 1xPBS. Cells were counterstained with 1:1000 Hoechst 33342. GFP-488 and Hoechst-350 fluorescence signals were acquired using an Opera Phenix Plus High-Content Screening System. The fluorescence signals were obtained from 15 fields of three wells and two plates. Fluorescence intensity levels were analyzed based on nuclear/area detection using the Hoechst-350 channel, and the 488 mean intensity levels within this area as the principal read out. Data analysis was performed at a single-cell level. Images were analyzed using the Harmony High-Content Imaging and Analysis Software, version 4.9.

---

**Extended Data Fig. 1.**
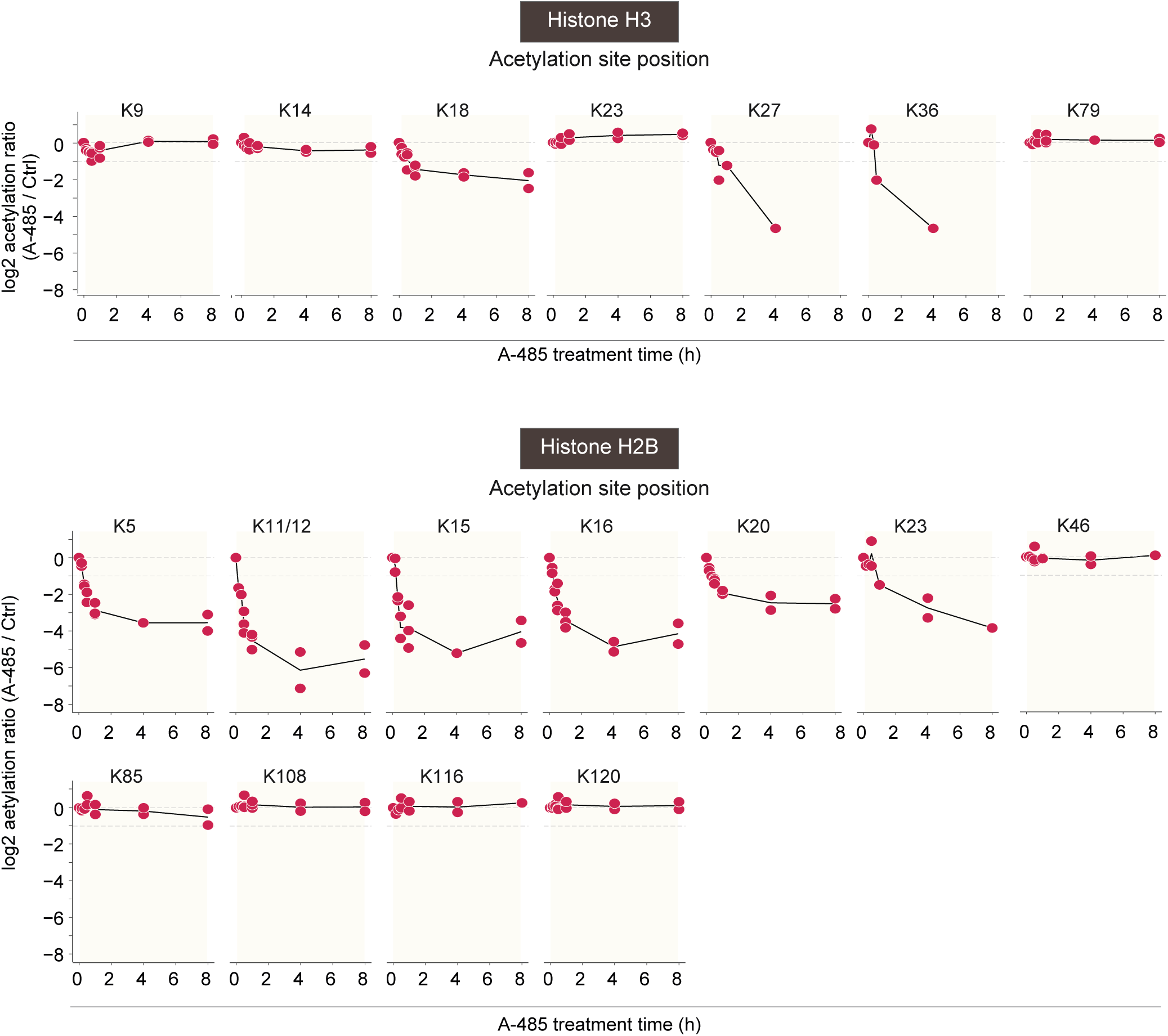
H3K27ac and H2BNTac are rapidly deacetylated after CBP/p300 inhibition. Deacetylation kinetics of histone H3 and H2B sites after CBP/p300 inhibition in mouse embryonic fibroblasts. Histone H3 and H2B acetylation site time-course data were re-analyzed from Weinert et al.^18^.

**Extended Data Fig. 2.**
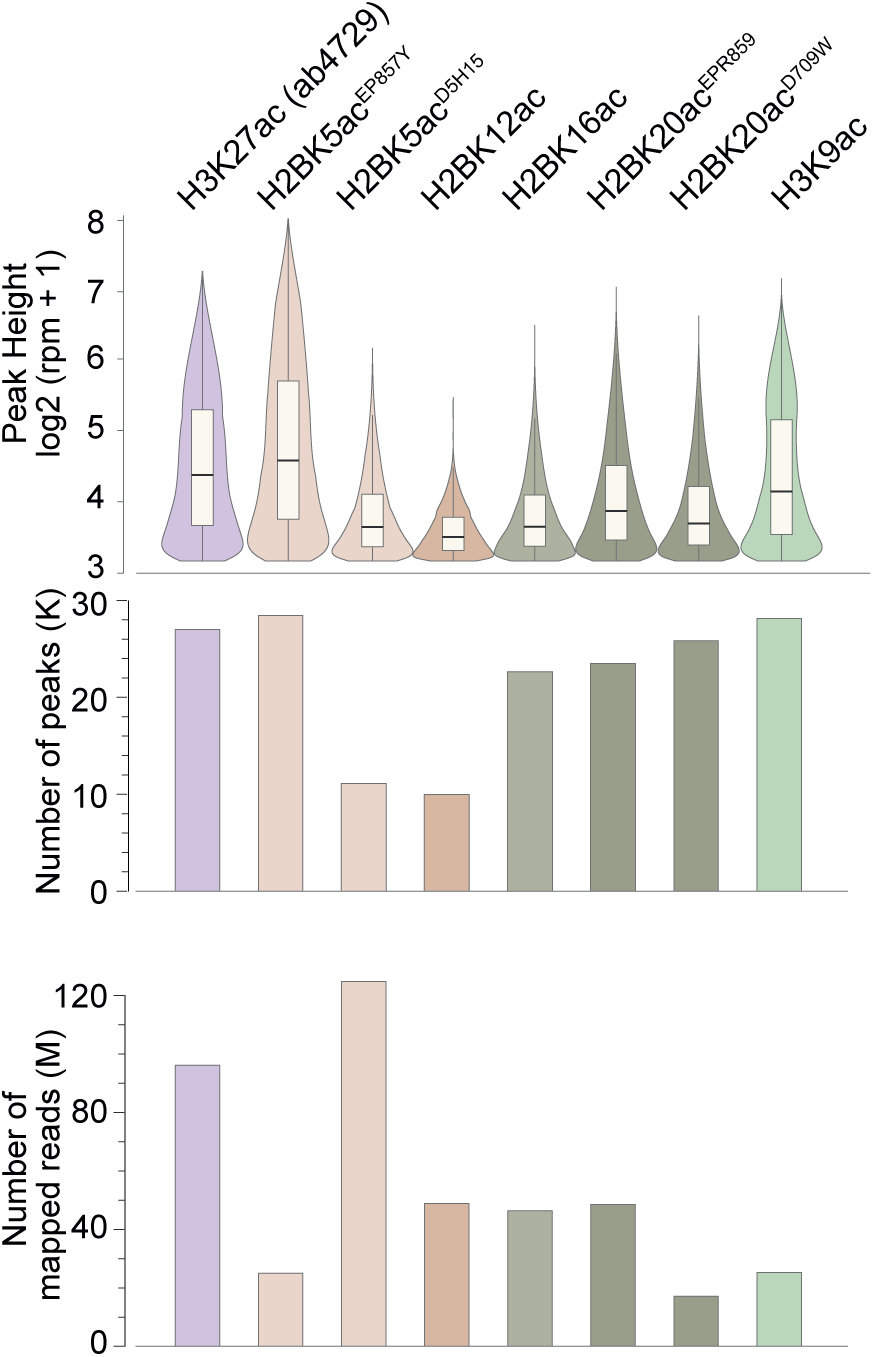
A summary of ChIP-seq analyses. The upper panel shows a comparison of H3K27ac, H3K9ac, and H2BNTac site ChIP signal intensity, the middle panel shows the number of identified peaks, and the lower panel shows the number of mapped read counts. Note that the number of identified peaks does not directly scale with the number of mapped reads. Instead, the number of identified peaks is better reflected in peak height; H2BK5ac^D5H15^ and H2BK12ac show the lowest peak height and the lowest number of identified peaks. Lower H2BK5ac^D5H15^ and H2BK12ac peak height may reflect weaker enrichment efficiency of the used antibodies and/or lower abundance of these marks.

**Extended Data Fig. 3.**
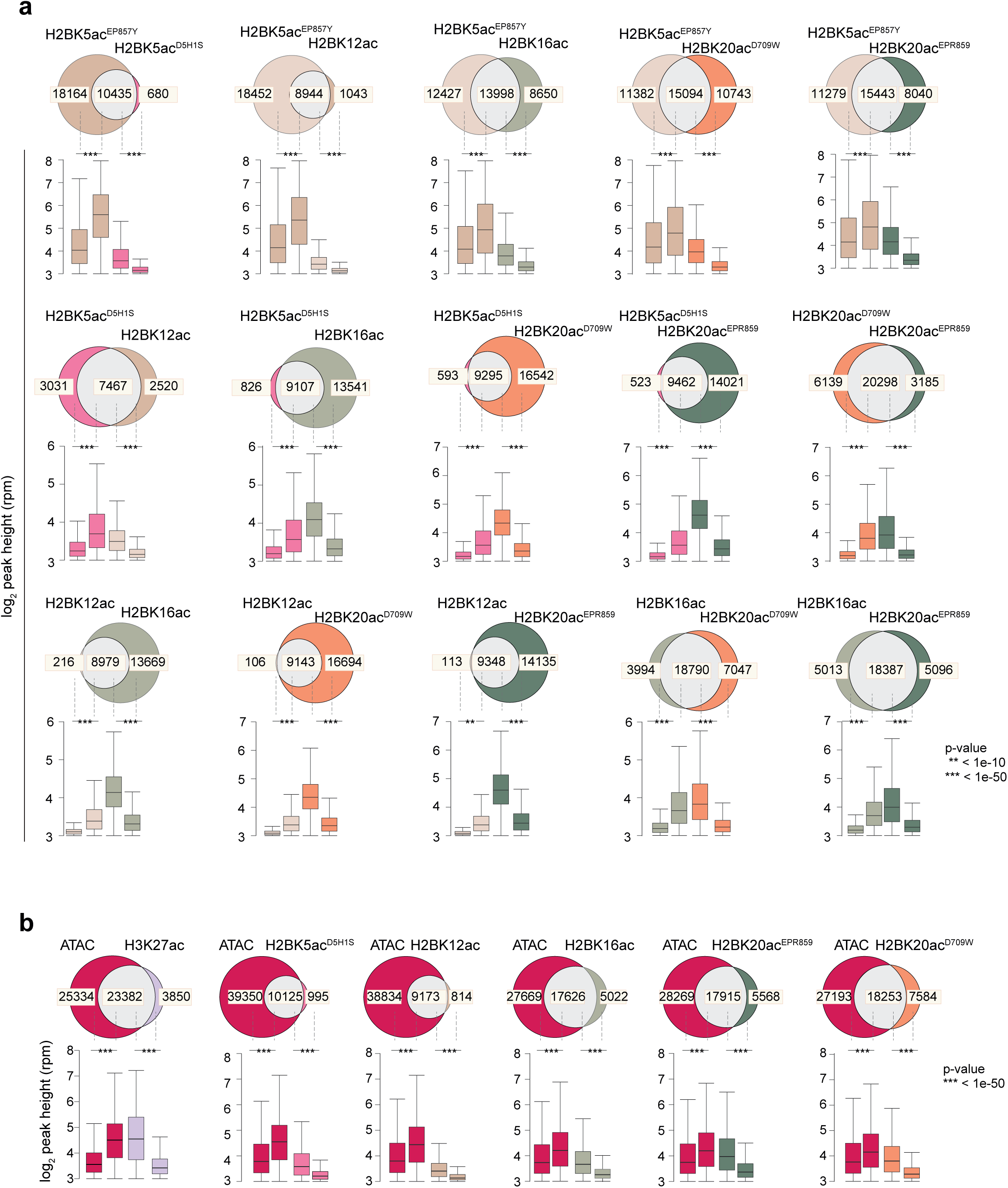
H2BNTac sites occupy the same genomic regions. **a,** Venn diagrams show the overlap of ChIP-seq peaks between the indicated H2BNTac marks. Below each Venn diagram, box plots show the distribution of peak height for the overlapping and non-overlapping peaks for each of the compared acetylation marks. Due to difficulty in peak calling and variation in peak width, a peak in one dataset can overlap with more than one peaks in another dataset or vice versa, which leads to a slightly different number of overlapping peaks between H2BNTac peaks. In this case, a higher peak number is shown as a representative overlapping peak number. The box plots display the median, upper and lower quartiles, and whiskers show 1.5× interquartile range (IQR). **b,** Overlap of H3K27ac and H2BNTac peaks with ATAC-seq peaks in mESC. Bar charts, below Venn diagrams, show the peak height of the indicated histone marks or ATAC-seq. The box plots display the median, upper and lower quartiles, and whiskers show 1.5× interquartile range (IQR). Statistical significance determined by Mann–Whitney U test.

**Extended Data Fig. 4.**
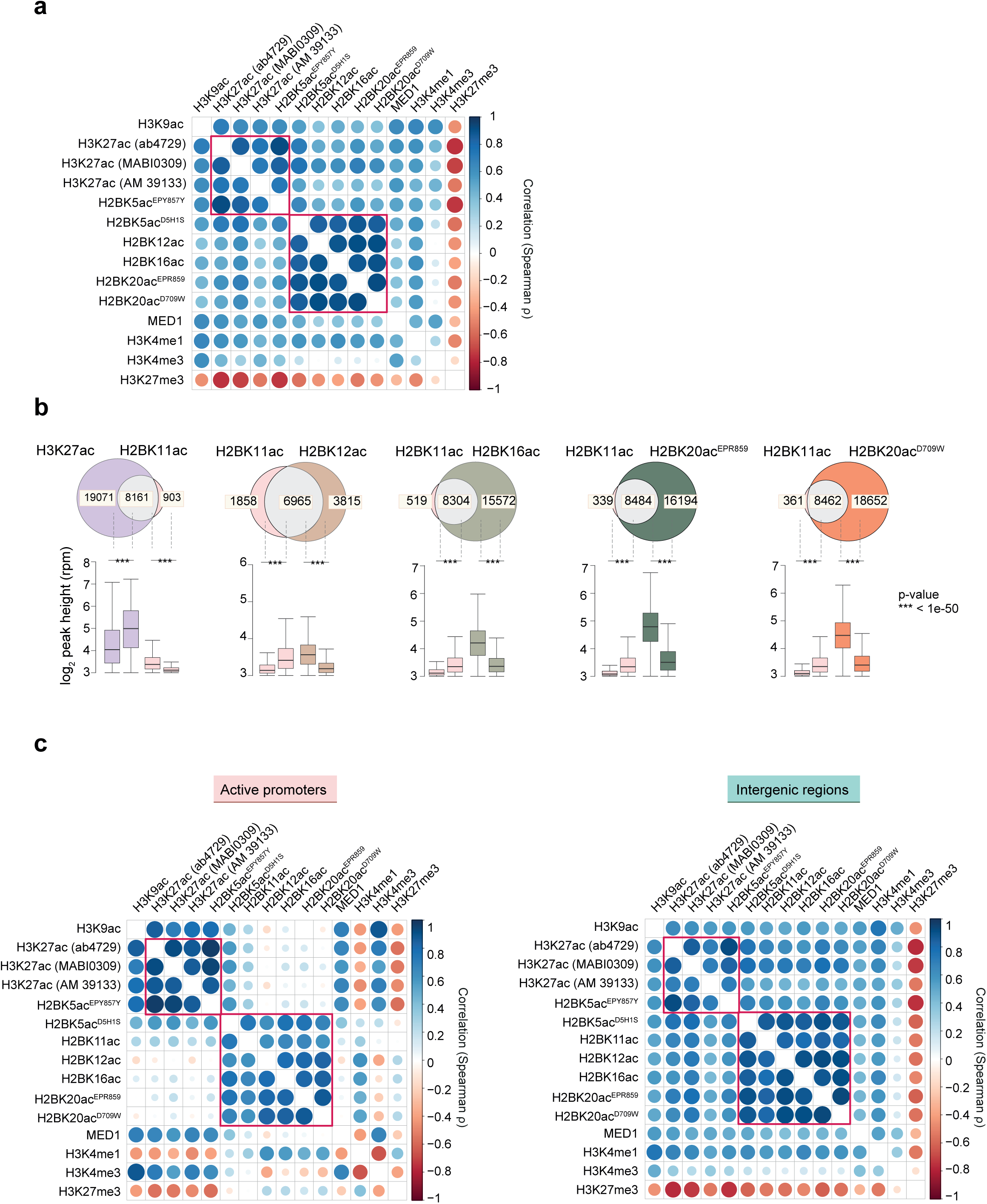
H3K27ac and H2BNTac distinctly correlate with other chromatin marks. **a,** Genome-wide correlation among the indicated chromatin marks. Correlation (Spearman ρ) is calculated between the indicated marks using a 2kb window in peak regions. H2BNTac strongly correlates with each other and shows a variable positive correlation with H3K4me1, H3K27ac, H3K9ac, and MED1. Notably, H2BNTac very poorly correlates with H3K4me3 and negatively correlates with H3K27me3. **b,** H2BK11ac overlaps with other H2BNTac sites. Shown is the overlap of H2BK11ac^+^ regions with the indicated H3K27ac and H2BNTac marks. Statistical significance determined by Mann–Whitney U test. **c,** Genome-wide correlation among the indicated chromatin marks. Correlation is determined as described above in a. H2BK11ac occupancy resembles that of other H2BNTac sites, and confirms the distinct occupancy profiles of H2BNTac and H3K27ac. As a note, the H2BK11ac antibody became available after most of the other experiments and analyses were completed. Because of this late addition of these data and limited coverage of H2B acetylated regions by this antibody, H2BK11ac was not included in other comparisons.

**Extended Data Fig. 5.**
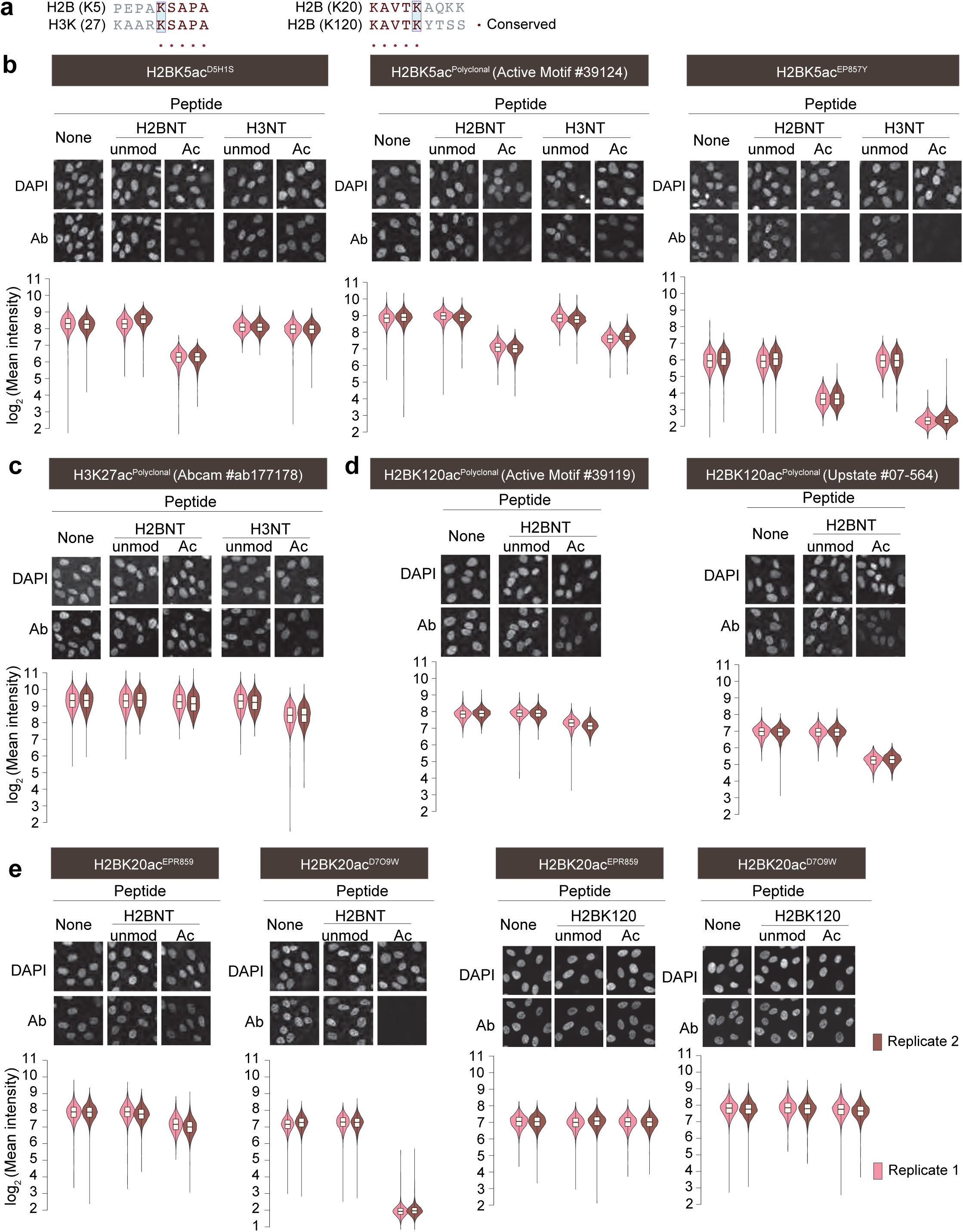
Previously used H2BK5ac and H2BK120ac ChIP antibodies cross-react with other histone acetylation sites. **a,** The sequence similarity of amino acids flanking H2BK5 and H3K27, and H2BK20 and H2BK120. Acetylated lysine is positioned in the middle and marked with a box. Identical amino acids are marked with dots. **b-e,** Analysis of H2BK5ac (b), H3K27ac (c), H2BK120ac (d), and H2BK20ac (e) antibody specificities. Cells were immunostained with the stated antibodies in the absence, or presence of the indicated unmodified (Unmod) or acetylated (Ac) peptides. Shown are the representative immunofluorescence images. Violin plots show the distribution of immunofluorescence signals for the indicated antibodies and treatment conditions. For each condition, 3,000 cells were quantified. Data are from 2 independent biological replicates. The box plots show the median, interquartile range (IQR), and whiskers show 1.5x IQR. Unmod (unmodified peptide), Ac (acetylated peptide), Ab (antibody), H2BNT (H2B N-terminus peptide), and H3NT (H3 N-terminus peptide). Peptide sequences used for antibody specificity testing are provided in Supplemental Table 2.

**Extended Data Fig. 6.**
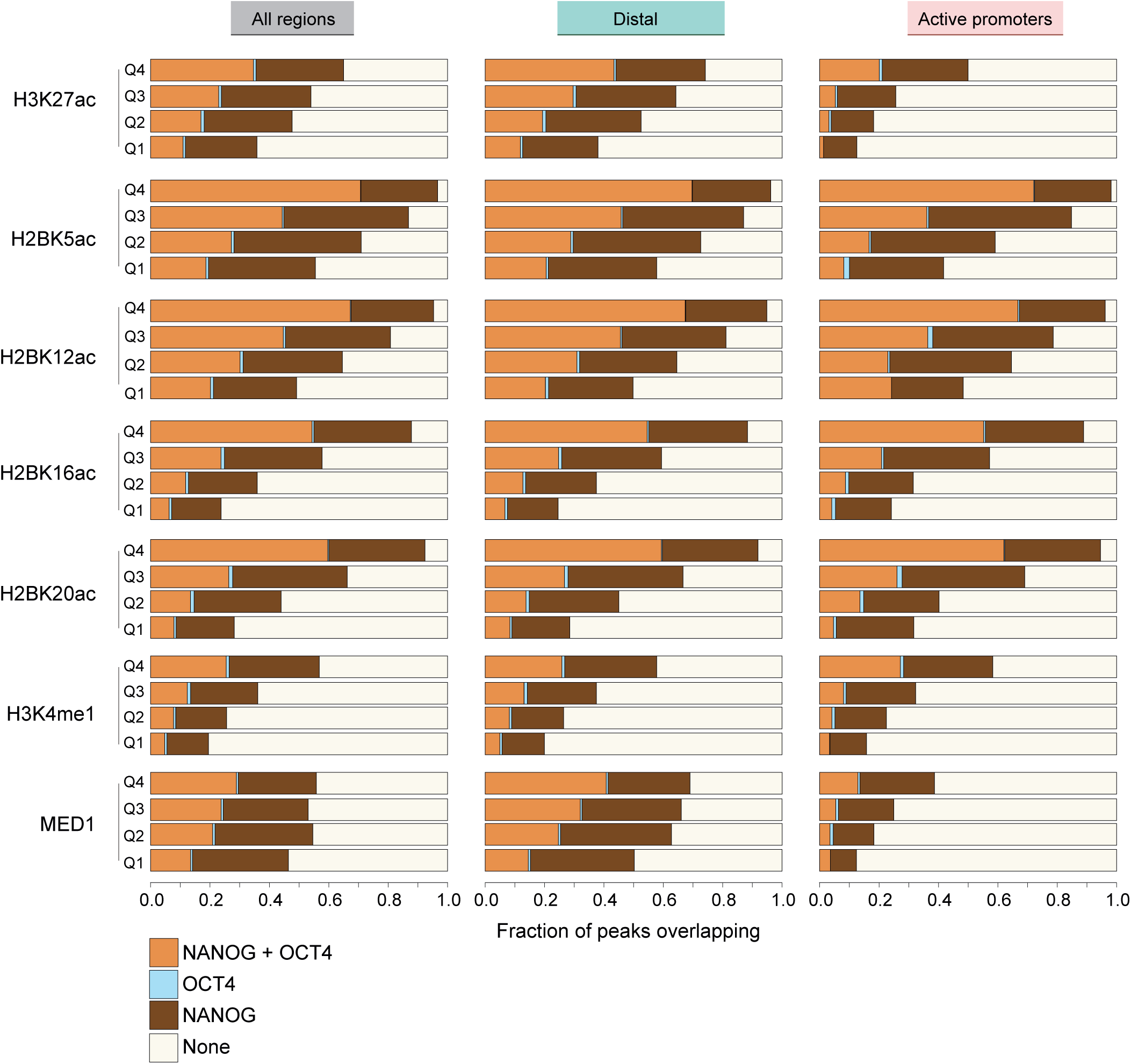
H2BNTac^+^ regions are occupied by NANOG and OCT4. Bar charts showing the fraction of H2BNTac, H3K27ac, H3K4me1, and MED1 regions bound by NANOG and/or OCT4. Based on ChIP signal intensity, H2BNTac, H3K27ac, H3K4me1, and MED1 regions were grouped into quartiles, and within each group, the fraction of regions overlapping with NANOG and/or OCT4 is shown. Co-occupancy of the indicated chromatin marks and OCT4 and/or NANOG was analyzed in the indicated genomic regions. All, all peaks; Active promoter, peaks mapping to ±1kb of active promoters (TPM ≥2 and marked with H3K4me3); Intergenic, peaks occurring outside promoter regions.

**Extended Data Fig. 7.**
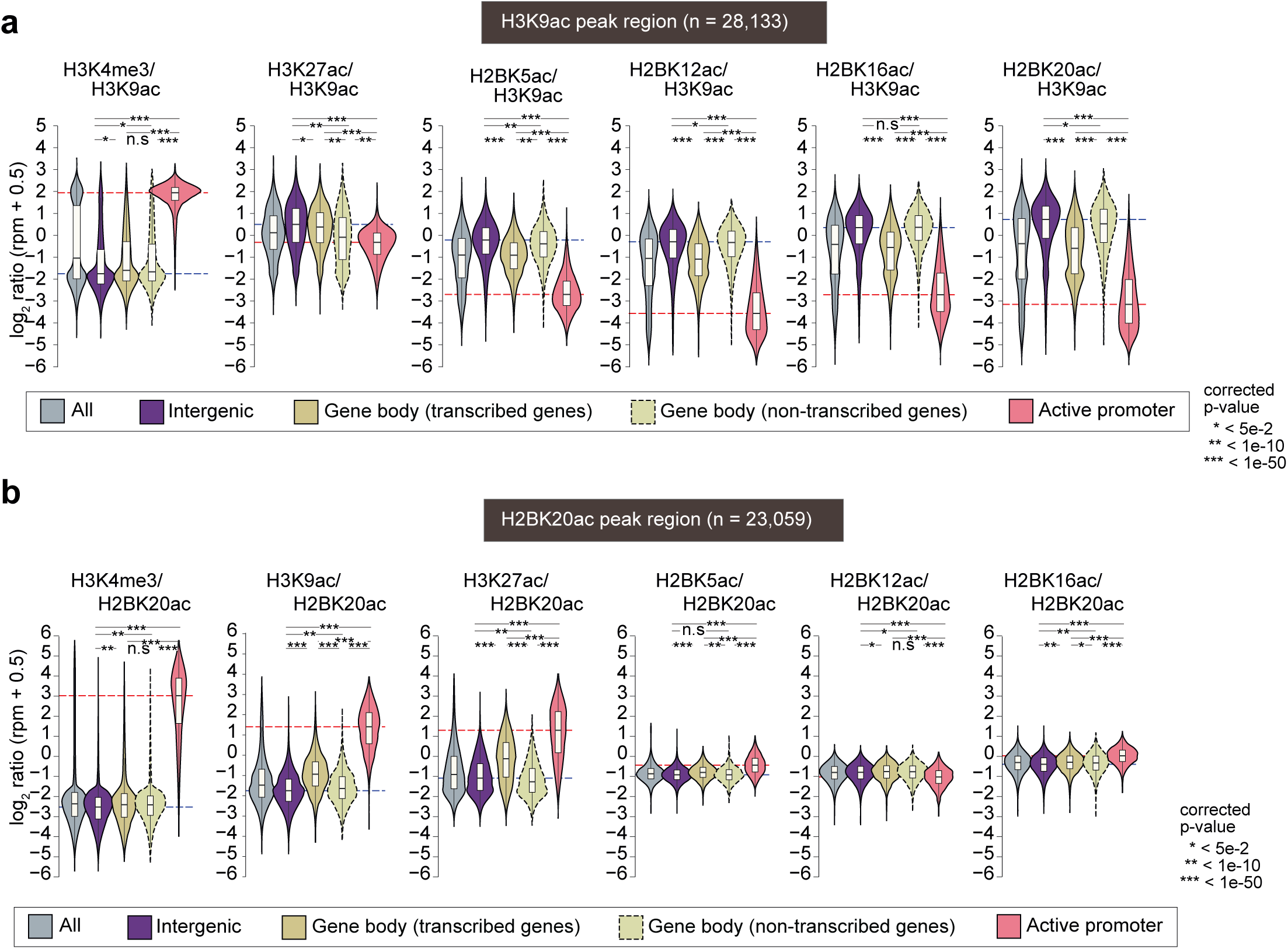
H2BNTac separates candidate enhancers and active promoters. a-b, Shown are the relative ChIP-seq signal intensities of the indicated chromatin marks at H3K9ac occupied regions (a) or H2BK20ac regions (b) in mESC. Active promoters (**±**1 kb from TSS) are defined as promoters of actively transcribed genes in mESC (TPM ≥2) and marked with H3K4me3. ChIP-seq peaks are grouped into the following categories: All, all peaks; Intergenic, peaks occurring outside promoters and gene body; Gene body (transcribed genes), peaks occurring within transcribed gene body; Gene body (non-transcribed genes), peaks occurring within the non-transcribed gene body. The dotted lines indicate the median ratio of the indicated chromatin marks at intergenic regions (blue) and active promoters (red). Statistical significance determined by Mann–Whitney U test.

**Extended Data Fig. 8.**
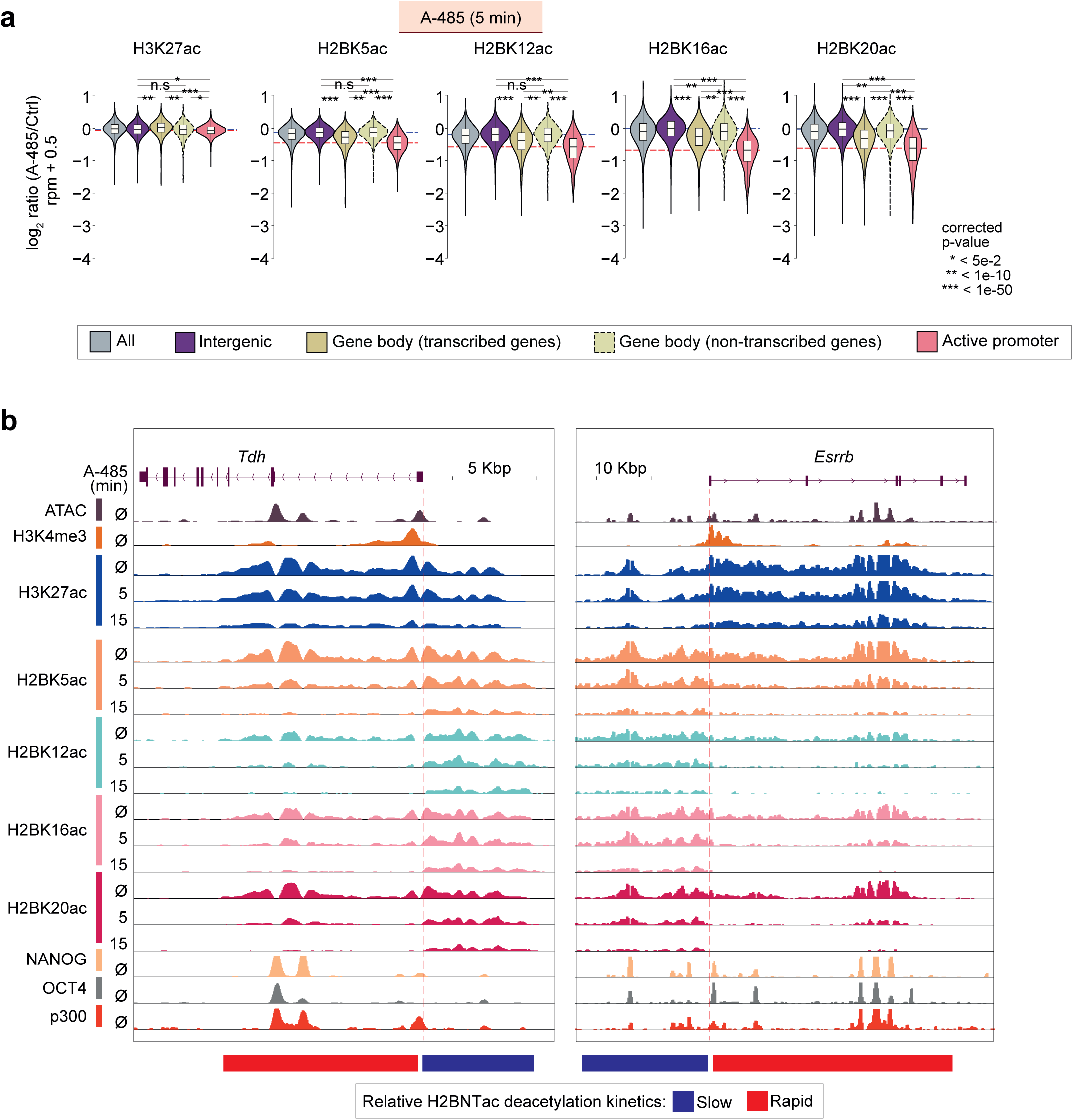
H2BK20ac is rapidly removed in actively transcribed promoters and gene body regions. **a,** CBP/p300 inhibition causes H2BNTac deacetylation within 5 minutes. Shown the fold-change in H3K27ac and H2BNTac site ChIP signal in untreated and A-485 treated (5 min) cells. Different genomic regions were classified as defined in Extended Data Fig. 7. Dotted lines indicate the median ratio at intergenic and active promoter regions. Statistical significance determined by Mann–Whitney U test. **b,** Genome browser view of Tdh and Essrb loci showing rapid removal of H2BNTac in actively transcribed gene body regions after CBP/p300 inhibition. As compared to intergenic regions, H2BNTac is more rapidly downregulated in enhancers occurring within transcribed gene body regions. In contrast, H3K27ac is similarly decreased at active promoters and candidate enhancers located in intergenic or gene body regions. The dotted lines demarcate intergenic regions (blue) and active promoters (red).

**Extended Data Fig. 9.**
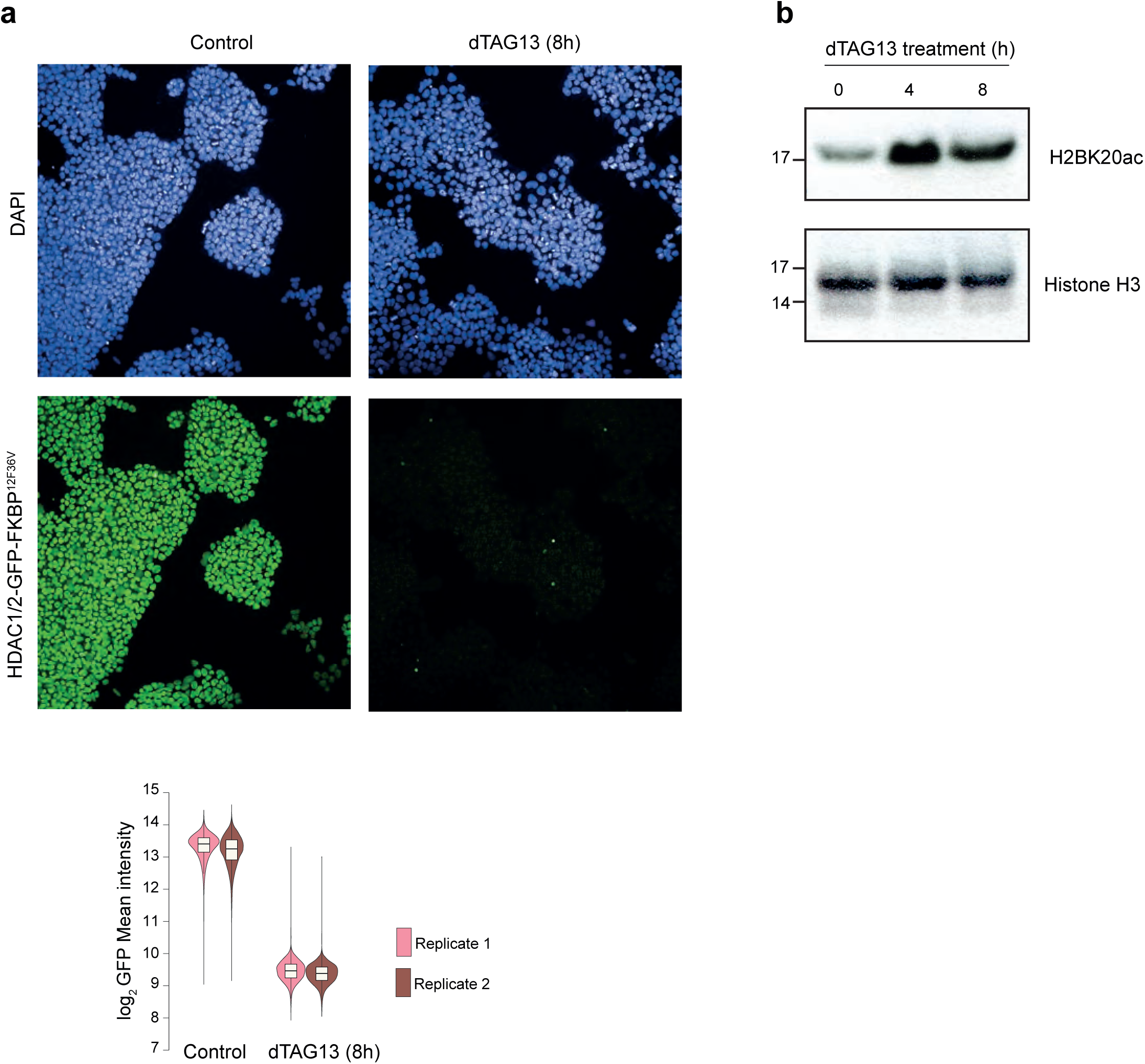
HDAC1/2 deacetylates H2BNTac. **a,** Endogenous HDAC1/2 were fused with GFP-FKBP12^F36V^ and acutely depleted by treating cells with dTAG13 (8h). Depletion of HDAC1/2-GFP-FKBP12^F36V^ was confirmed by analyzing GFP expression by microscopy. The representative images show loss of GFP signal in dTAG13 treated HDAC1/2-GFP-FK-BP12^F36V^ cells. Violin plots show the distribution of GFP signals for the indicated treatment conditions. For each condition, >29,000 cells were quantified. Data are from 2 independent biological replicates. The box plots show the median, interquartile range (IQR), and whiskers show 1.5x IQR. **b,** Immunoblot showing acetylation of H2BK20 in control and HDAC1/2 depleted mESC (n=2).

**Extended Data Fig. 10.**
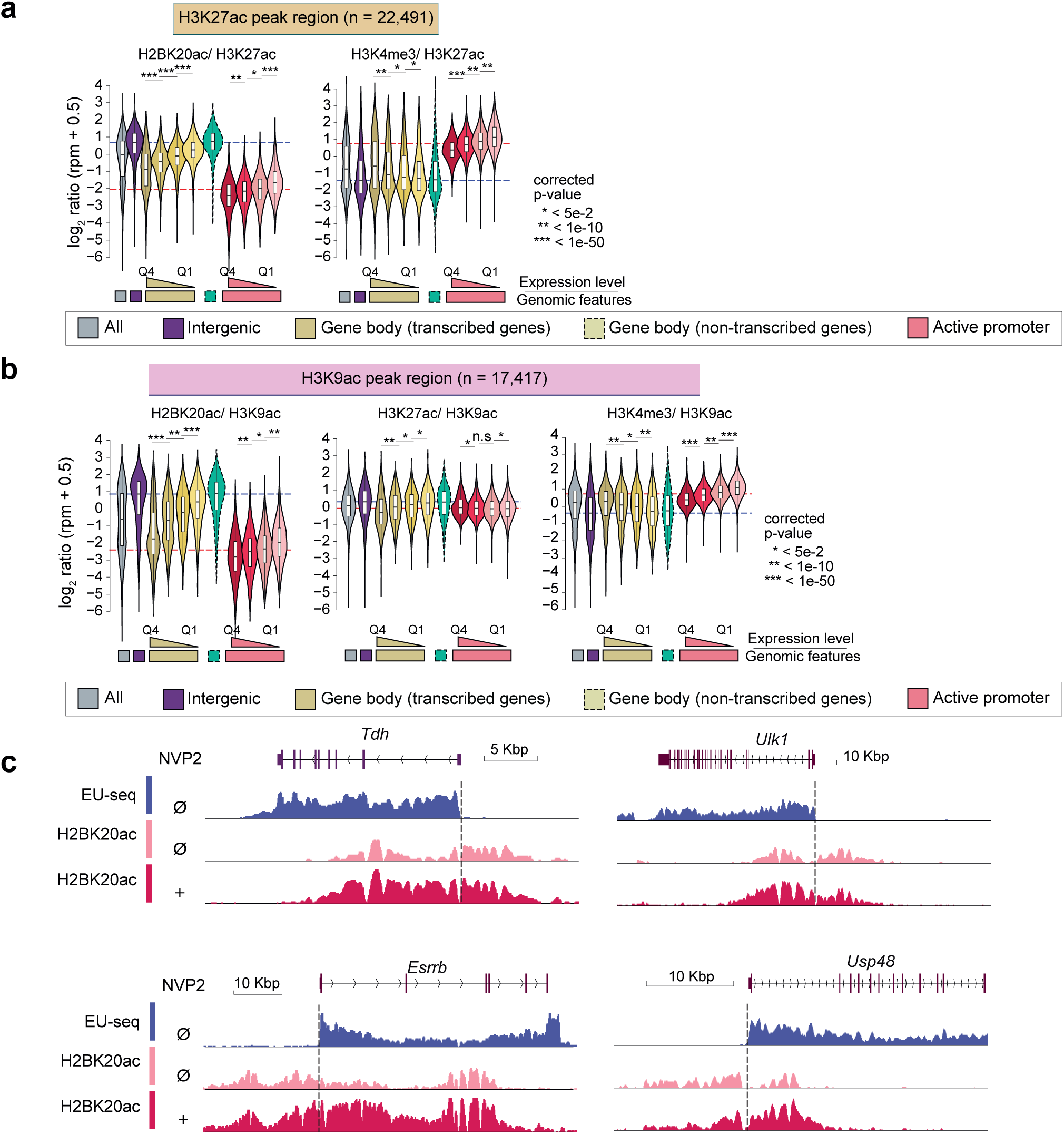
H2BNTac signal is lower in promoters and actively transcribed regions, and transcription inhibition increases H2BNTac. **a-b,** Relative enrichment of H2BK20ac and the indicated histone marks in H3K27ac regions (a) and H3K9ac regions (b) in K562. Actively transcribed genes in K562 were grouped into quartiles based on gene expression. Shown is the ratio of the indicated marks in the specified genomic regions. Note that H2BK20ac/H3K9ac and H2BK20ac/H3K27ac ratio in promoters and actively transcribed gene body regions are inversely related to gene expression. All, all peaks; intergenic, peaks occurring outside promoters and gene body; gene body (transcribed genes), peaks occurring within transcribed gene body; gene body (non-transcribed genes), peaks occurring within the non-transcribed gene body. Active promoters (±1 kb from TSS) are defined as promoters that are linked to actively transcribed genes in K562 (RNA-seq, TPM ≥2) and marked with H3K4me3. The dotted lines indicate the median ratio of the indicated chromatin marks at intergenic regions (blue) and active promoters (red). **c,** Representative genome browser tracks showing transcription inhibitor (NVP2)-induced preferential increase in H2BK20ac in actively transcribed regions. H2BK20ac ChIP-seq was performed in mESC treated without or with NVP2 (2h). The dotted line demarcates intergenic actively transcribed regions from upstream intergenic regions. Statistical significance determined by Mann–Whitney U test.

**Extended Data Fig. 11.**
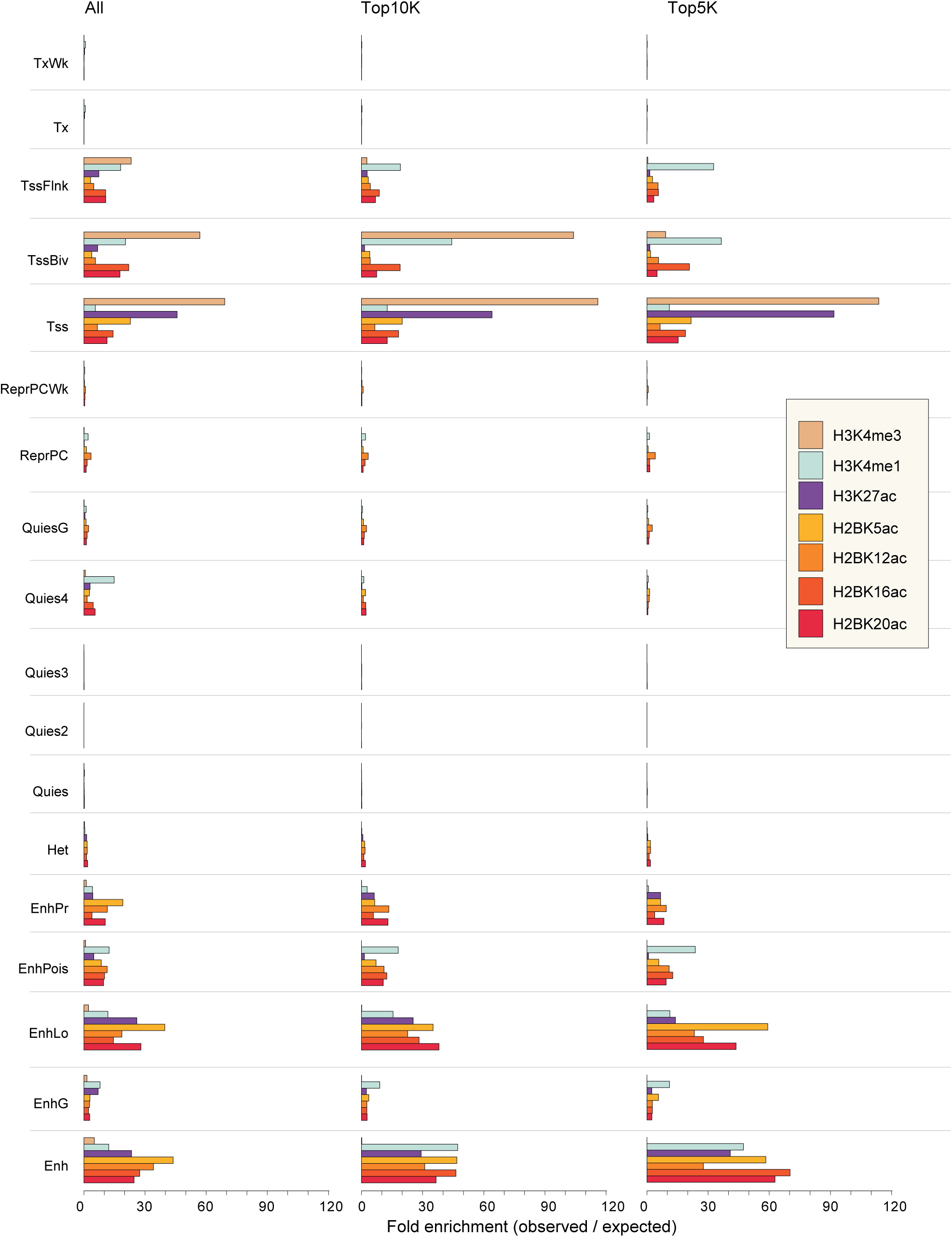
Enrichment of H2BNTac in ChromHMM defined chromatin states in mESC. Shown is the enrichment of the specified histone marks in the indicated ChromHMM-defined chromatin states. Top ten thousand (Top10K) and top five thousand (Top 5K) peaks were chosen based on the peak heights of the specified chromatin marks. mESC Chrom-HMM annotations were obtained from van der Velde et al.^61^.

**Extended Data Fig. 12.**
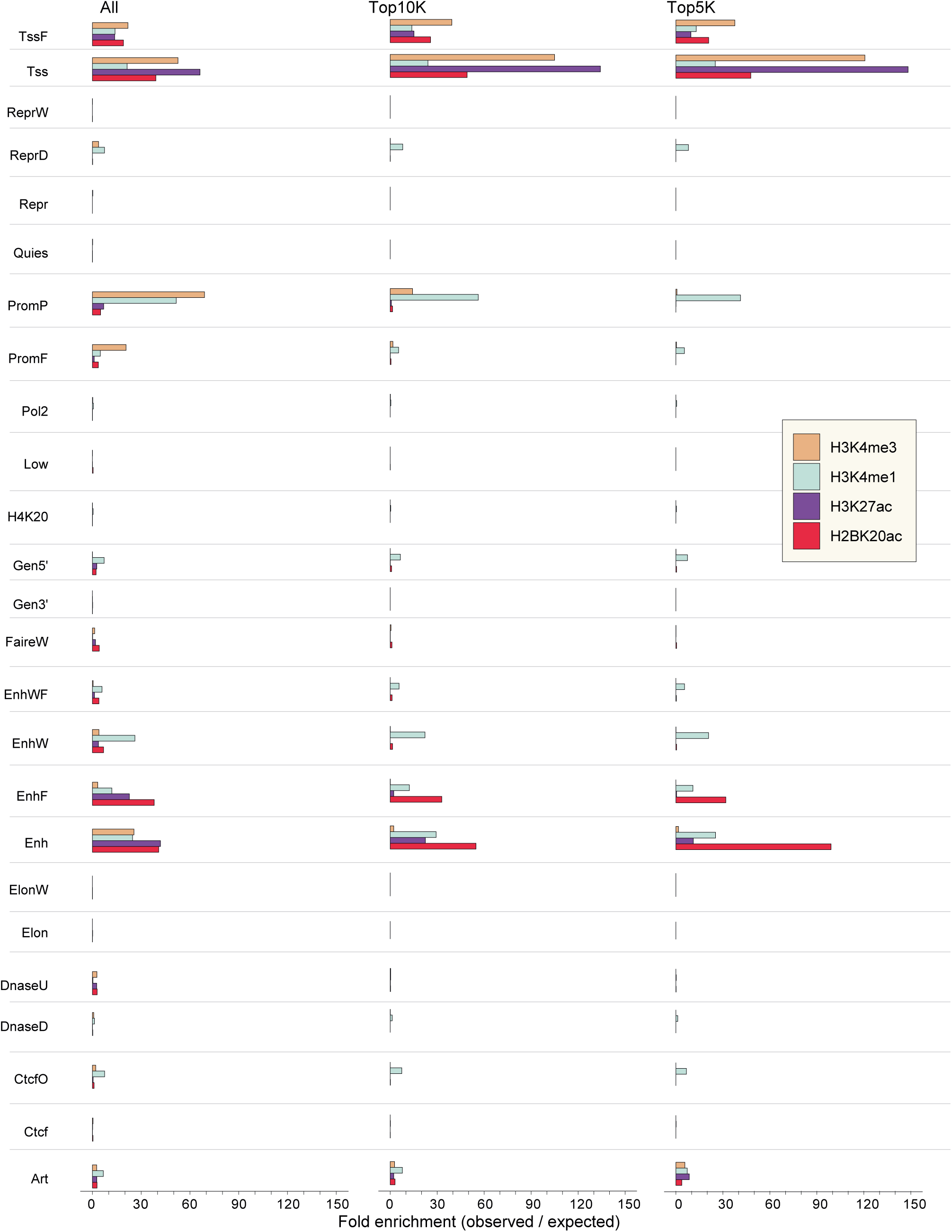
Enrichment of H2BNTac in ChromHMM defined chromatin states in K562 cells. Shown is the enrichment of the specified histone marks in the indicated ChromHMM-defined chromatin states. Top ten thousand (Top10K) and top five thousand (Top 5K) peaks were chosen based on the peak heights of the specified chromatin marks. K562 Chrom-HMM annotations were downloaded from (http://hgdownload.cse.ucsc.edu/goldenpath/hg19/encodeDCC/w-gEncodeAwgSegmentation)^34,62^

**Extended Data Fig. 13.**
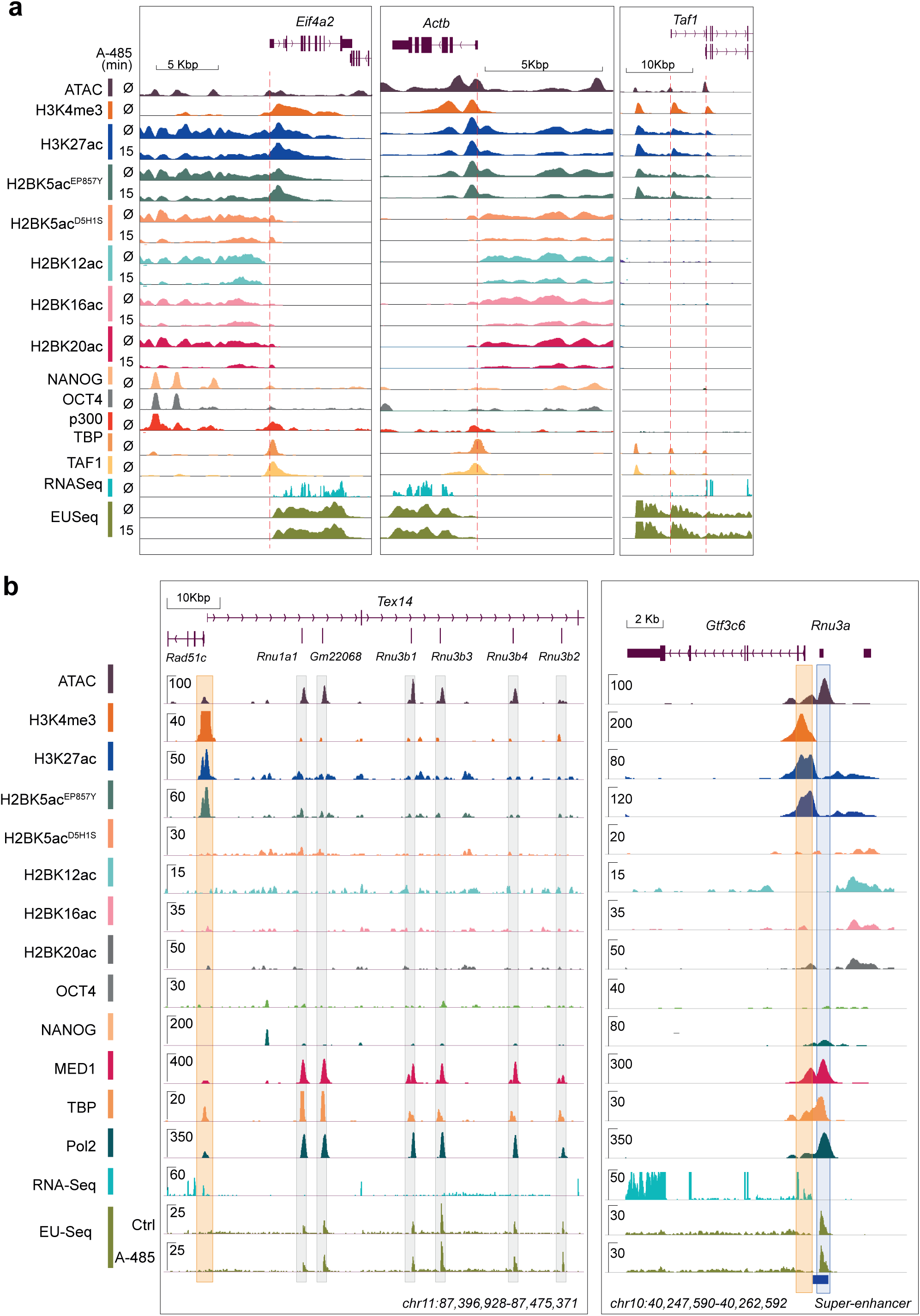
H2BNTac distinguishes candidate enhancers from protein-coding and snoRNA-coding gene promoters. **a,** H2BNTac distinguishes candidate enhancers from proximally occurring promoters of the housekeeping genes Actb and Eif4a2. The dotted line demarcates intergenic H2BK20ac^+^ candidate enhancer regions from the gene promoters. Taf1 upstream H3K27ac^+^ peak lacks H2BNTac but is marked with H3K4me3, identifying it as a putative promoter. This is consistent with the EU-seq signal in this region. The dotted line demarcates intergenic and the currently annotated Taf1 TSS. **b,** A genome browser view of Rad51c and Tex14 proximal (left) and Gtf3c6 proximal (right) regions. *Rad51c, Tex14,* and *Gtf3c6* promoters are shadowed in orange, and *Gtf3c6* upstream candidate super-enhancer^63^ and the six regions that were previously interpreted as *Rad51c*-proximal enhancers^35^, are shadowed in light blue and light grey, respectively. The candidate enhancers, indicated by shadowed boxes, are marked with a high level of ATAC-seq, MED1, TBP, and Pol2, but have little undetectable H3K4me3 and H2BNTac. While *Rad51c, Tex14, and Gtf3c6* expression are detected in RNA-seq, the candidate enhancer regions lack a detectable RNA-seq signal. Instead, they show a very high level of nascent RNA transcription, which remains unaffected by the A-485 treatment. As indicated, a closer inspection of the expanded genome annotation reveals all seven regions as snoRNA genes.

**Extended Data Fig. 14.**
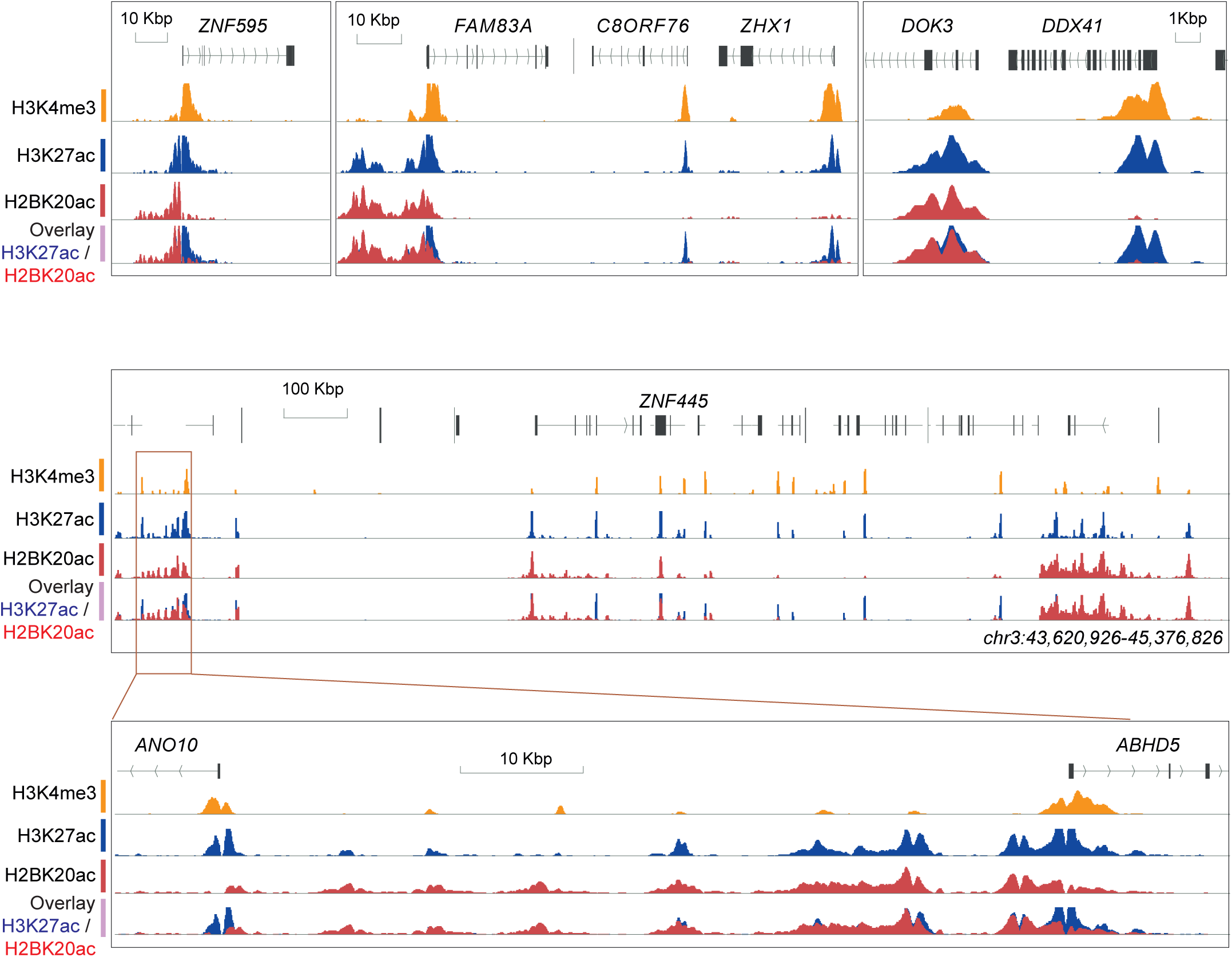
H2BK20ac distinguishes candidate enhancers from active promoters in human cells. Genome browser view of representative regions showing distinct occupancy profiles of H3K4me3, H3K27ac, and H2BK20ac in K562 cells. Note that H2BK20ac and H3K27ac show distinct enrichment in promoters. Some promoters are marked by both H2BK20ac and H3K27ac, whereas others are only marked by H3K27ac. Distal regions, marked with low H3K4me3, are similarly marked by H2BK20ac and H3K27ac.

**Extended Data Fig. 15.**
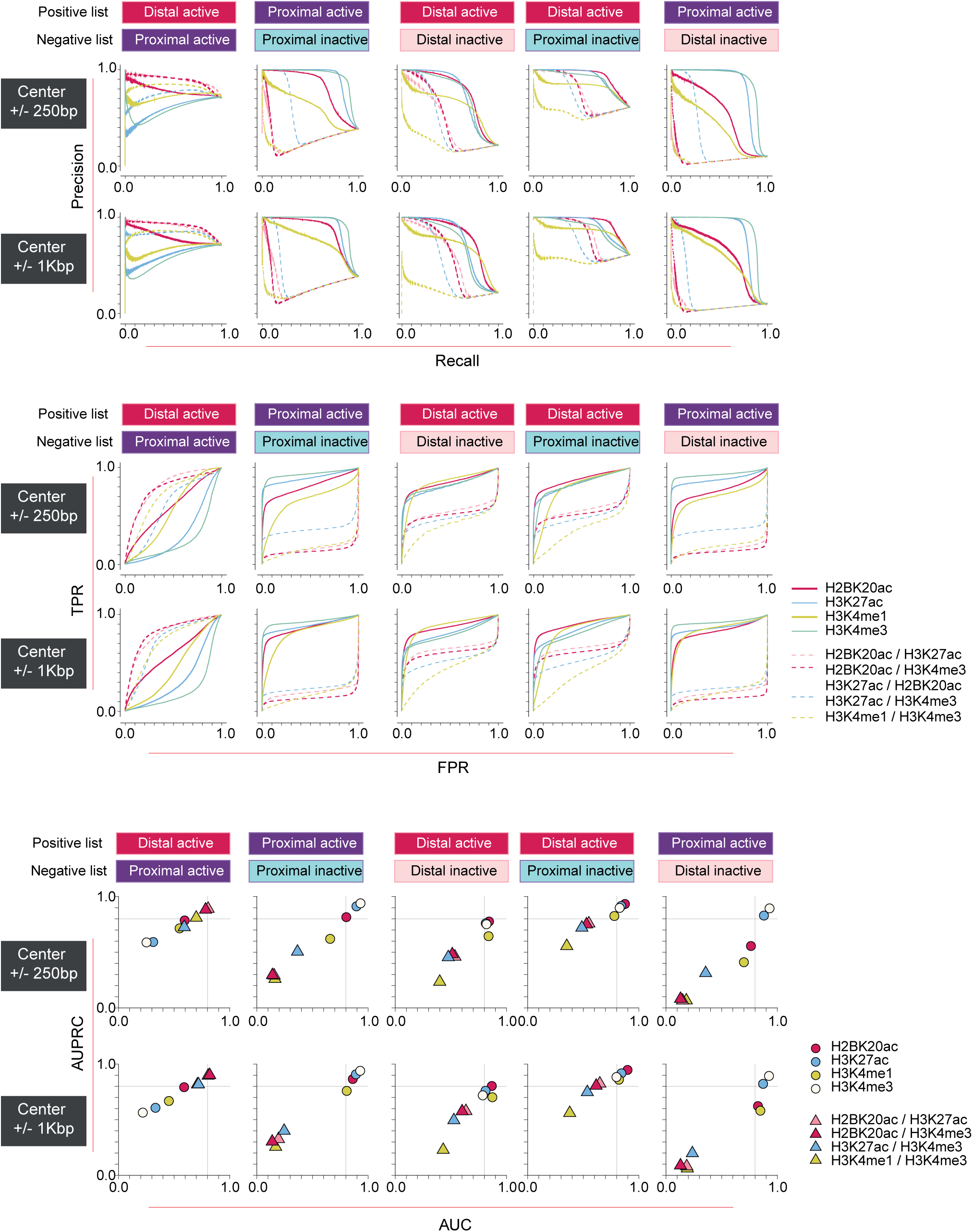
H2BNTac, H3K27ac, H3K4me3, and H3K4me1 discriminate transcriptionally active and inactive regions. Benchmarking chromatin marks for discriminating promoters and candidate enhancers. The discriminatory potential of different histone marks and histone mark ratios was analyzed by generating receiver operating characteristics (ROC) and precision-recall curves (PRC). True positive rate (TPR), false-positive rate (FPR), precision and recall were calculated under the different thresholds of peak enrichment or peak enrichment ratios. Transcriptionally active and inactive regions in K562 were defined based on RNA expression evidence in K562 and non-K562 cells and tissues^14^. PINTS regions were further classified as proximal or distal as defined by Yao et.al.^14^ (see Methods). We computed the ChIP-seq signal in +/-250 and 1,000bp of the PINTS region center. PINTS regions were grouped into four categories. Proximal active: proximal PINTS regions that are actively transcribed in K562 cells. Proximal inactive: proximal PINTS regions that are active in other cell types but not in K562, and occurred >2kb away from any K562 proximal active PINTS peaks. Distal active: distal PINTS regions that are actively transcribed in K562 cells. Distal inactive: distal PINTS regions that are active in other cell types but not in K562. Distal inactive regions that occurred in proximity to active PINTS regions were excluded from the analyses (see Methods).

**Extended Data Fig. 16.**
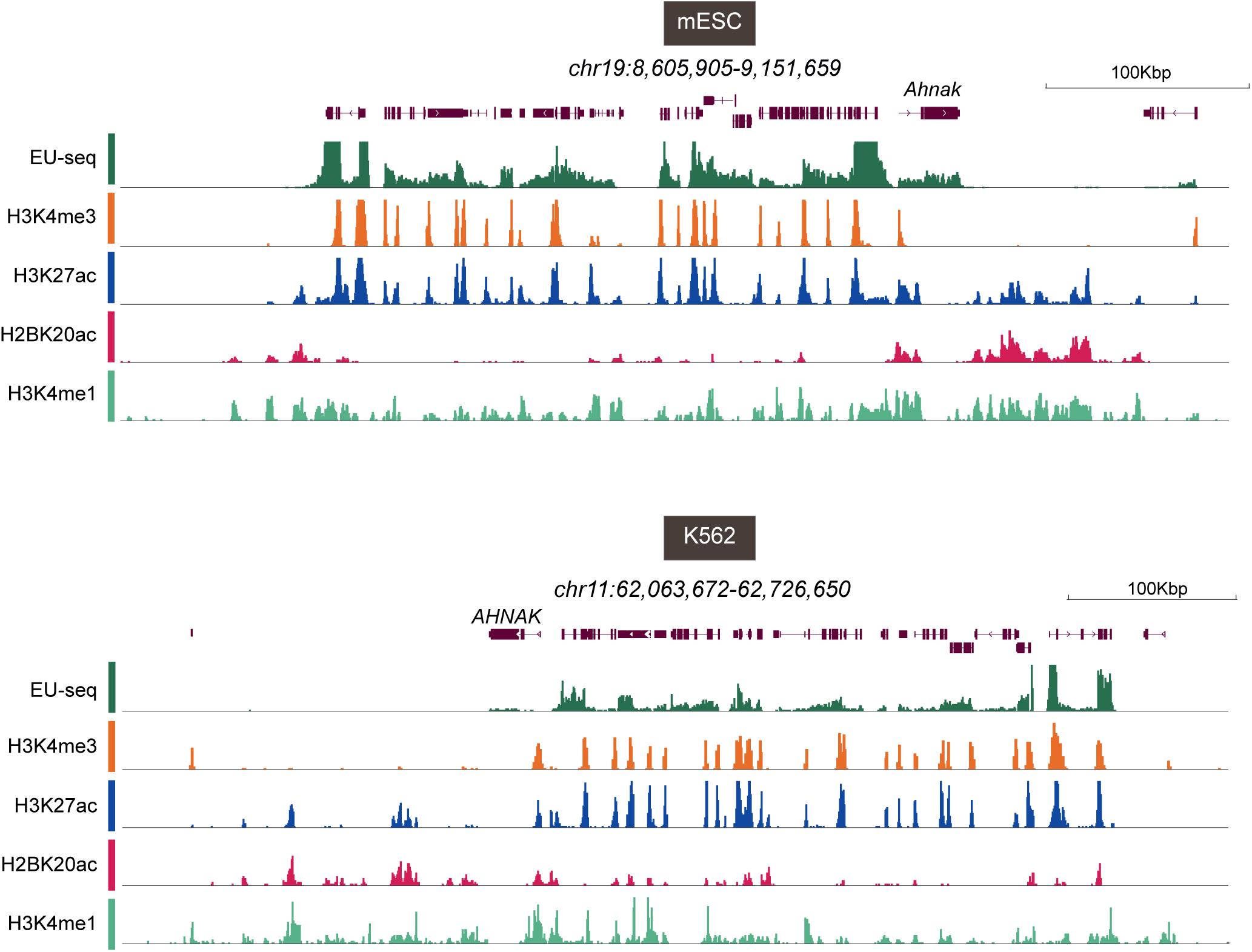
H2BNTac, H3K27ac, H3K4me3, and H3K4me1 show distinct genomic occupancy patterns. A Representative genomic regions in mESC (upper panel) and K562 (lower panel) showing different occupancy patterns of H3K27ac, H2BK20ac, H3K4me3, and H3K4me1 in promoters and candidate enhancer regions. Most of the active promoters, marked with a high level of H3K27ac and H3K4me3, show a low level of H2BK20ac. Candidate enhancers, near the Ahnak and AHNAK regions, display low H3K4me3 and are co-marked with a similar level of H2BK20ac and H3K27ac.

**Extended Data Fig. 17.**
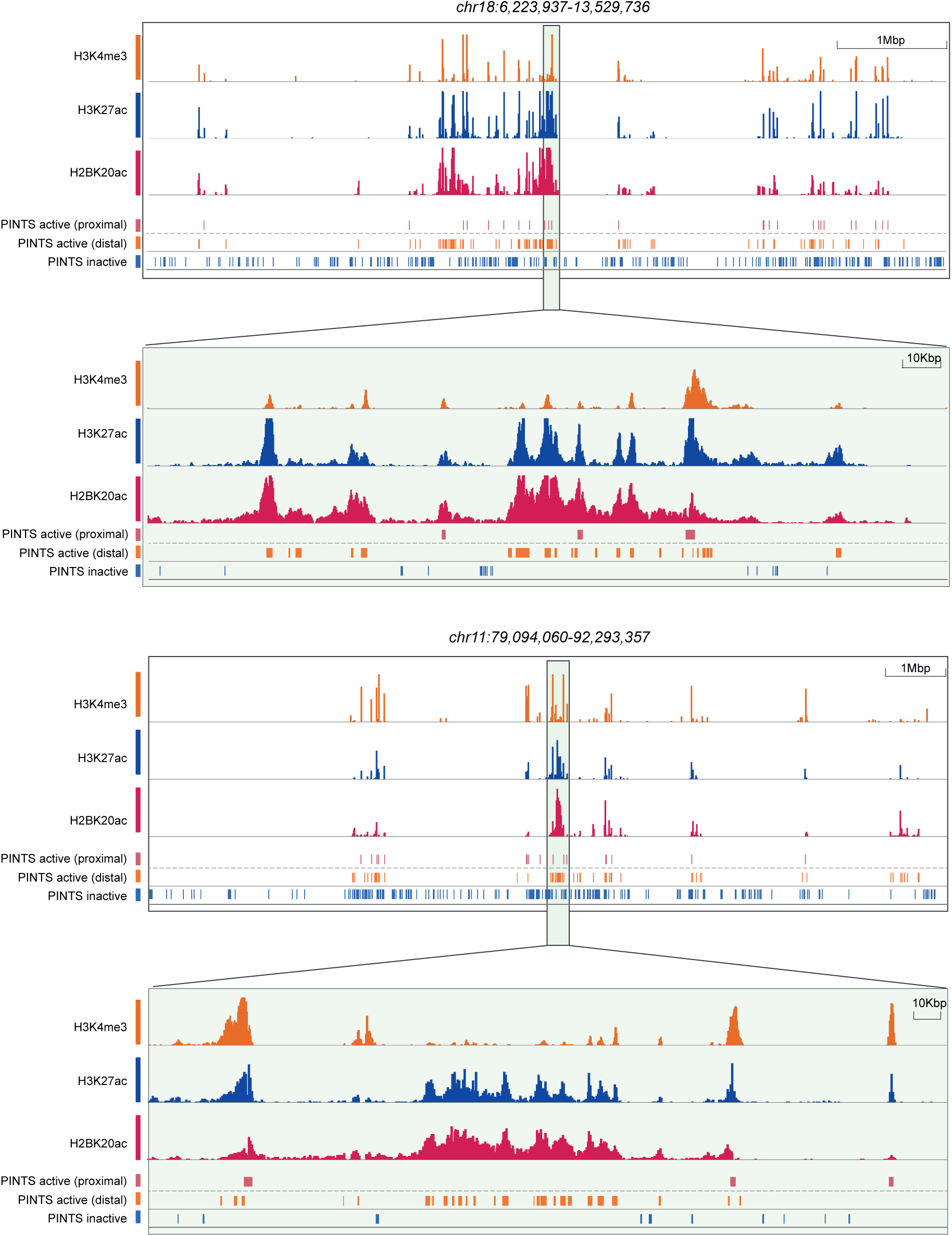
H2BNTac and H3K27ac prominently mark active PINTS regions. Representative genome browser tracks showing the occurrence of H2BNTac and H3K27ac in active and inactive PINTS regions. Two representative regions, and their zoom-in, are shown. The shown regions include hundreds of active and inactive PINTS regions. Of note, a vast majority of active PINTS regions are marked with H2BNTac and H3K27ac, whereas most inactive PINTS regions are either devoid of these acetylation marks or have a much lower level of acetylation. The distinct overlap of acetylation marks with active and inactive PINTS regions is clearer in the zoom-in regions showing a high overlap of acetylation marks with active PINTS regions, and low or lack of acetylation in inactive PINTS regions.

**Extended Data Fig. 18.**
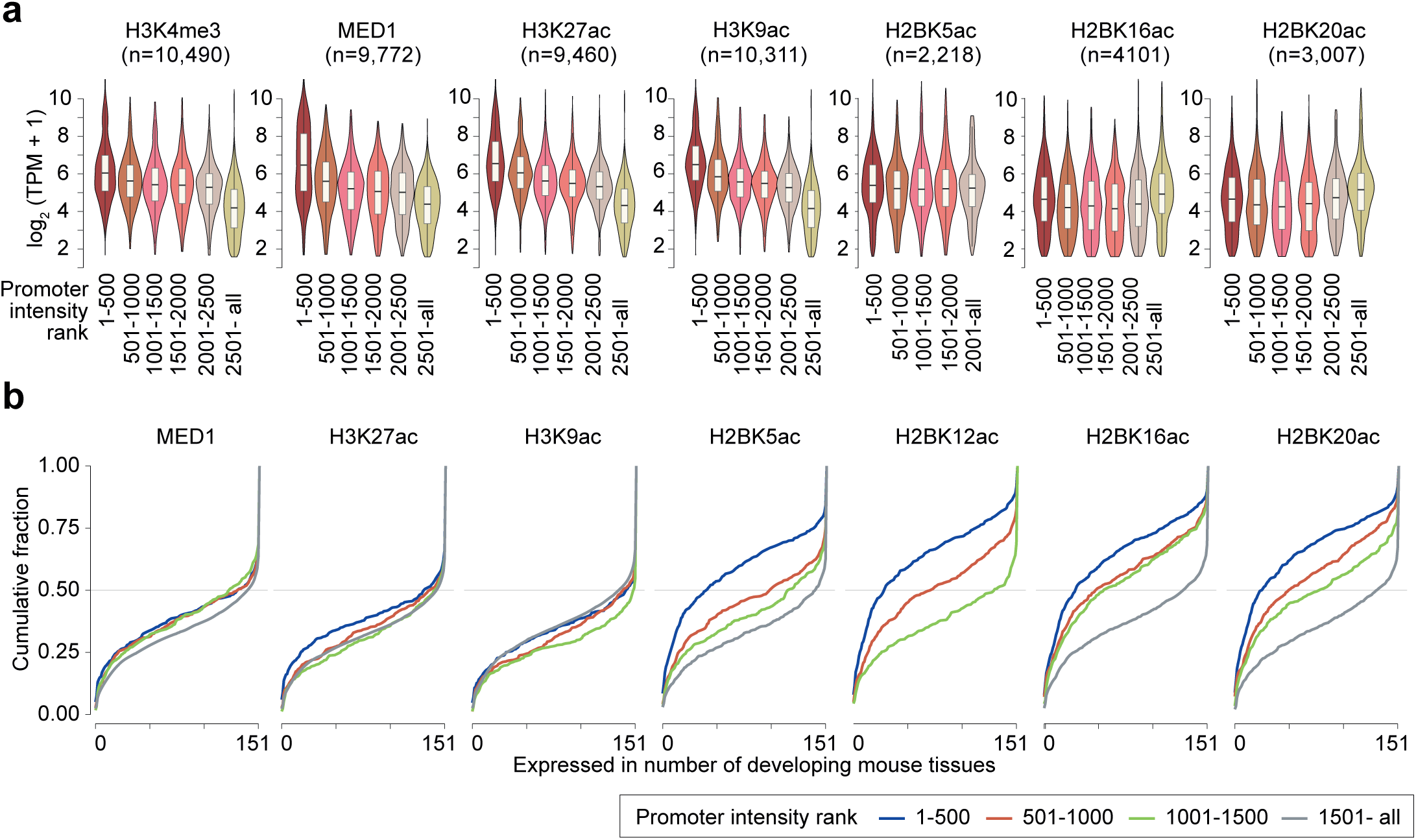
Promoter H2BNTac signal poorly associates with gene expression levels but indicates cell-type-specificity of genes. **a,** The indicated chromatin marks were rank-ordered based on their ChIP-seq signal intensity in promoters (± 1kb from TSS) of actively transcribed genes in mESC. Shown are gene expressions (TPM +1) within the indicated abundance ranges. **b,** H2BNTac preferentially marks cell-type-specific gene promoters. The indicated chromatin marks were rank-ordered and grouped into different intensity ranges, as described in a. Cell-type-specificity was determined within the indicated abundance ranges. Shown is a cumulative fraction of genes in each group plotted against genes expressed in 151 FANTOM5 CAGE profiles of mouse tissues ^24^.

**Extended Data Fig. 19.**
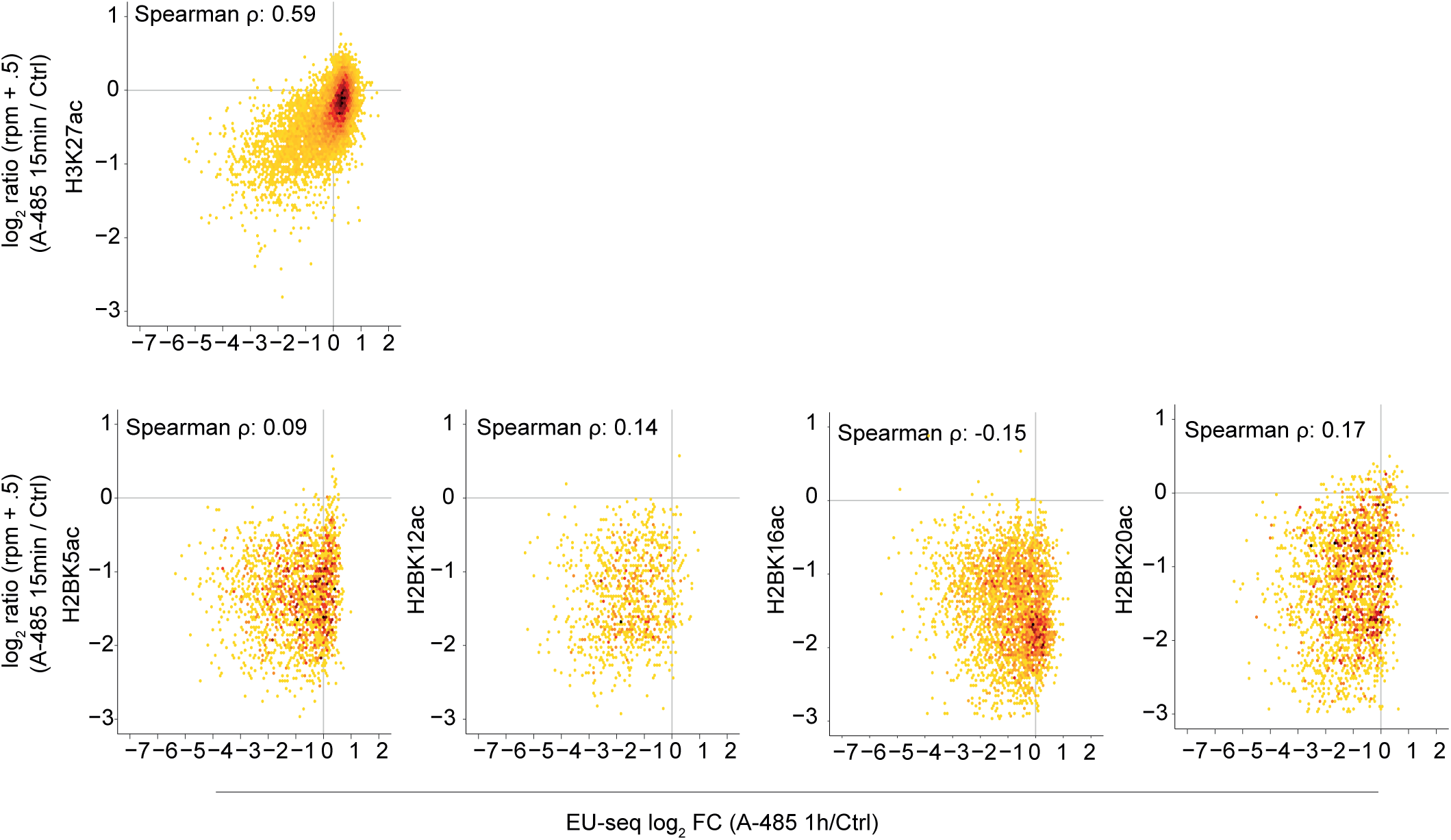
CBP/p300 inhibitor-induced downregulation of promoter-associated H3K27ac correlates with gene downregulation. Correlation (Spearman r) between A-485-induced changes in promoter-associated H3K27ac or H2BNTac site intensity and A-485-induced gene downregulation. mESC were treated without or with A-485 (15 min) and change in H3K27ac and H2BNTac signal was determined using ChIP-seq. H3K27ac and H2BNTac intensity were quantified within the ±1kb region from TSS. A-485-induced changes in nascent transcription were determined after 1 h of A-485 treatment ^21^.

**Extended Data Fig. 20.**
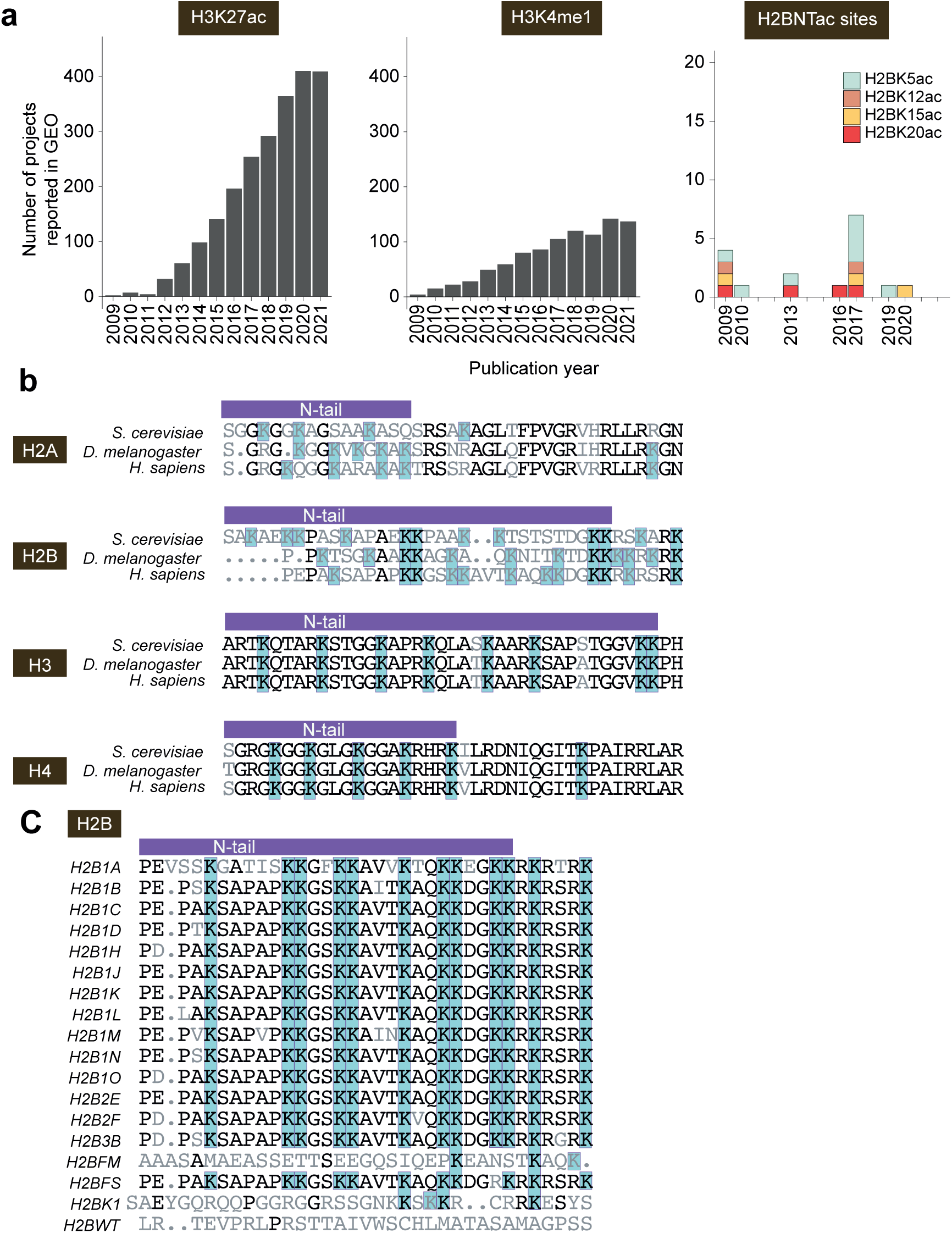
H3K27ac is extensively profiled, whereas H2BNTac is vastly understudied and the H2BNT sequence is poorly conserved among eukaryotes. **a,** Plotted is the number of H3K27ac, H3K4me1, and H2BNTac site genome-wide mapping projects deposited in the GEO repository from 2009 till 2021. Note that, over the past decade, the number of H3K27ac genome-wide profiling projects reported in GEO has linearly increased year to year. In contrast, the number of deposited H2BNTac projects has not changed over the years, showing that H2BNTac remains understudied. H2BK5ac is profiled more often than other H2BNTac marks, but our analyses indicate that the antibodies used for previously published H2BK5ac ChIP-seq analyses cross-react with H3K27ac (**Extended Data** Fig. 5). Note that the figure shows the number of projects (not the number of individual ChIP-seq experiments) that included genome-wide mapping data for the indicated marks. **b,** The sequence similarity of core histone N-termini among the indicated species. As compared to histones H3 and H4 N-termini, the histone H2B N-terminus shows high sequence divergence. **c,** The sequence similarity of H2BNT among different human histone H2B isoforms.

## References

1. Calo, E. & Wysocka, J. Modification of enhancer chromatin: what, how, and why? Mol Cell 49, 825–837, doi:10.1016/j.molcel.2013.01.038 (2013).

2. Shlyueva, D., Stampfel, G. & Stark, A. Transcriptional enhancers: from properties to genome-wide predictions. Nat Rev Genet 15, 272–286, doi:10.1038/nrg3682 (2014).

3. Andersson, R. & Sandelin, A. Determinants of enhancer and promoter activities of regulatory elements. Nat Rev Genet 21, 71–87, doi:10.1038/s41576-019-0173-8 (2020).

4. Bernstein, B. E. et al. The NIH Roadmap Epigenomics Mapping Consortium. Nat Biotechnol 28, 1045–1048, doi:10.1038/nbt1010-1045 (2010).

5. Stunnenberg, H. G., International Human Epigenome, C. & Hirst, M. The International Human Epigenome Consortium: A Blueprint for Scientific Collaboration and Discovery. Cell 167, 1145–1149, doi:10.1016/j.cell.2016.11.007 (2016).

6. Consortium, E. P. et al. Expanded encyclopaedias of DNA elements in the human and mouse genomes. Nature 583, 699–710, doi:10.1038/s41586-020-2493-4 (2020).

7. Taylor, G. C., Eskeland, R., Hekimoglu-Balkan, B., Pradeepa, M. M. & Bickmore, W. A. H4K16 acetylation marks active genes and enhancers of embryonic stem cells, but does not alter chromatin compaction. Genome Res 23, 2053–2065, doi:10.1101/gr.155028.113 (2013).

8. Pradeepa, M. M. et al. Histone H3 globular domain acetylation identifies a new class of enhancers. Nat Genet 48, 681–686, doi:10.1038/ng.3550 (2016).

9. Kumar, V. et al. Comprehensive benchmarking reveals H2BK20 acetylation as a distinctive signature of cell-state-specific enhancers and promoters. Genome Res 26, 612–623, doi:10.1101/gr.201038.115 (2016).

10. Regadas, I. et al. A unique histone 3 lysine 14 chromatin signature underlies tissue- specific gene regulation. Mol Cell 81, 1766–1780 e1710, doi:10.1016/j.molcel.2021.01.041 (2021).

11. Creyghton, M. P. et al. Histone H3K27ac separates active from poised enhancers and predicts developmental state. Proc Natl Acad Sci U S A 107, 21931–21936, doi:10.1073/pnas.1016071107 (2010).

12. Field, A. & Adelman, K. Evaluating Enhancer Function and Transcription. Annu Rev Biochem 89, 213–234, doi:10.1146/annurev-biochem-011420-095916 (2020).

13. Andersson, R. et al. An atlas of active enhancers across human cell types and tissues. Nature 507, 455–461, doi:10.1038/nature12787 (2014).

14. Yao, L. et al. A comparison of experimental assays and analytical methods for genome-wide identification of active enhancers. Nat Biotechnol, doi:10.1038/s41587-022-01211-7 (2022).

15. Barakat, T. S. et al. Functional Dissection of the Enhancer Repertoire in Human Embryonic Stem Cells. Cell Stem Cell 23, 276–288 e278, doi:10.1016/j.stem.2018.06.014 (2018).

16. Gasperini, M., Tome, J. M. & Shendure, J. Towards a comprehensive catalogue of validated and target-linked human enhancers. Nat Rev Genet 21, 292–310, doi:10.1038/s41576-019-0209-0 (2020).

17. Galouzis, C. C. & Furlong, E. E. M. Regulating specificity in enhancer-promoter communication. Curr Opin Cell Biol 75, 102065, doi:10.1016/j.ceb.2022.01.010 (2022).

18. Weinert, B. T. et al. Time-Resolved Analysis Reveals Rapid Dynamics and Broad Scope of the CBP/p300 Acetylome. Cell 174, 231–244 e212, doi:10.1016/j.cell.2018.04.033 (2018).

19. Scholz, C. et al. Acetylation site specificities of lysine deacetylase inhibitors in human cells. Nat Biotechnol 33, 415–423, doi:10.1038/nbt.3130 (2015).

20. Hansen, B. K. et al. Analysis of human acetylation stoichiometry defines mechanistic constraints on protein regulation. Nat Commun 10, 1055, doi:10.1038/s41467-019-09024-0 (2019).

21. Narita, T. et al. Enhancers are activated by p300/CBP activity-dependent PIC assembly, RNAPII recruitment, and pause release. Mol Cell 81, 2166–2182 e2166, doi:10.1016/j.molcel.2021.03.008 (2021).

22. Wang, Z. et al. Combinatorial patterns of histone acetylations and methylations in the human genome. Nat Genet 40, 897–903, doi:10.1038/ng.154 (2008).

23. Rajagopal, N. et al. Distinct and predictive histone lysine acetylation patterns at promoters, enhancers, and gene bodies. G3 (Bethesda) 4, 2051-2063, doi:10.1534/g3.114.013565 (2014).

24. Lasko, L. M. et al. Discovery of a selective catalytic p300/CBP inhibitor that targets lineage-specific tumours. Nature 550, 128–132, doi:10.1038/nature24028 (2017).

25. Egelhofer, T. A. et al. An assessment of histone-modification antibody quality. Nat Struct Mol Biol 18, 91–93, doi:10.1038/nsmb.1972 (2011).

26. Heintzman, N. D. et al. Histone modifications at human enhancers reflect global cell-type-specific gene expression. Nature 459, 108–112, doi:10.1038/nature07829 (2009).

27. Whyte, W. A. et al. Master transcription factors and mediator establish super-enhancers at key cell identity genes. Cell 153, 307–319, doi:10.1016/j.cell.2013.03.035 (2013).

28. Chen, X. et al. Integration of external signaling pathways with the core transcriptional network in embryonic stem cells. Cell 133, 1106–1117, doi:10.1016/j.cell.2008.04.043 (2008).

29. Wang, Z. A. et al. Histone H2B Deacylation Selectivity: Exploring Chromatin’s Dark Matter with an Engineered Sortase. J Am Chem Soc 144, 3360–3364, doi:10.1021/jacs.1c13555 (2022).

30. Nabet, B. et al. The dTAG system for immediate and target-specific protein degradation. Nat Chem Biol 14, 431–441, doi:10.1038/s41589-018-0021-8 (2018).

31. Olson, C. M. et al. Pharmacological perturbation of CDK9 using selective CDK9 inhibition or degradation. Nat Chem Biol 14, 163–170, doi:10.1038/nchembio.2538 (2018).

32. Venkatesh, S. & Workman, J. L. Histone exchange, chromatin structure and the regulation of transcription. Nat Rev Mol Cell Biol 16, 178–189, doi:10.1038/nrm3941 (2015).

33. Yaakov, G., Jonas, F. & Barkai, N. Measurement of histone replacement dynamics with genetically encoded exchange timers in yeast. Nat Biotechnol 39, 1434–1443, doi:10.1038/s41587-021-00959-8 (2021).

34. Consortium, E. P. An integrated encyclopedia of DNA elements in the human genome. Nature 489, 57–74, doi:10.1038/nature11247 (2012).

35. Hua, P. et al. Defining genome architecture at base-pair resolution. Nature 595, 125–129, doi:10.1038/s41586-021-03639-4 (2021).

36. Peng, T. et al. STARR-seq identifies active, chromatin-masked, and dormant enhancers in pluripotent mouse embryonic stem cells. Genome Biol 21, 243, doi:10.1186/s13059-020-02156-3 (2020).

37. Visel, A. et al. ChIP-seq accurately predicts tissue-specific activity of enhancers. Nature 457, 854–858, doi:10.1038/nature07730 (2009).

38. Shen, Y. et al. A map of the cis-regulatory sequences in the mouse genome. Nature 488, 116–120, doi:10.1038/nature11243 (2012).

39. Heintzman, N. D. et al. Distinct and predictive chromatin signatures of transcriptional promoters and enhancers in the human genome. Nat Genet 39, 311–318, doi:10.1038/ng1966 (2007).

40. Kwasnieski, J. C., Fiore, C., Chaudhari, H. G. & Cohen, B. A. High-throughput functional testing of ENCODE segmentation predictions. Genome Res 24, 1595–1602, doi:10.1101/gr.173518.114 (2014).

41. Schoenfelder, S. & Fraser, P. Long-range enhancer-promoter contacts in gene expression control. Nat Rev Genet 20, 437–455, doi:10.1038/s41576-019-0128-0 (2019).

42. Karlic, R., Chung, H. R., Lasserre, J., Vlahovicek, K. & Vingron, M. Histone modification levels are predictive for gene expression. Proc Natl Acad Sci U S A 107, 2926–2931, doi:10.1073/pnas.0909344107 (2010).

43. Fulco, C. P. et al. Activity-by-contact model of enhancer-promoter regulation from thousands of CRISPR perturbations. Nat Genet 51, 1664–1669, doi:10.1038/s41588-019-0538-0 (2019).

44. Hsieh, T. S. et al. Resolving the 3D Landscape of Transcription-Linked Mammalian Chromatin Folding. Mol Cell 78, 539–553 e538, doi:10.1016/j.molcel.2020.03.002 (2020).

45. Krietenstein, N. et al. Ultrastructural Details of Mammalian Chromosome Architecture. Mol Cell 78, 554–565 e557, doi:10.1016/j.molcel.2020.03.003 (2020).

46. Halfon, M. S. Studying Transcriptional Enhancers: The Founder Fallacy, Validation Creep, and Other Biases. Trends Genet 35, 93–103, doi:10.1016/j.tig.2018.11.004 (2019).

47. Sahu, B. et al. Sequence determinants of human gene regulatory elements. Nat Genet 54, 283–294, doi:10.1038/s41588-021-01009-4 (2022).

48. Ing-Simmons, E. et al. Independence of chromatin conformation and gene regulation during Drosophila dorsoventral patterning. Nat Genet 53, 487–499, doi:10.1038/s41588-021-00799-x (2021).

49. Espinola, S. M. et al. Cis-regulatory chromatin loops arise before TADs and gene activation, and are independent of cell fate during early Drosophila development. Nat Genet 53, 477–486, doi:10.1038/s41588-021-00816-z (2021).

50. Struhl, K. Histone acetylation and transcriptional regulatory mechanisms. Genes Dev 12, 599–606, doi:10.1101/gad.12.5.599 (1998).

51. Cramer, P. Organization and regulation of gene transcription. Nature 573, 45–54, doi:10.1038/s41586-019-1517-4 (2019).

52. Filippakopoulos, P. & Knapp, S. The bromodomain interaction module. FEBS Lett 586, 2692–2704, doi:10.1016/j.febslet.2012.04.045 (2012).

53. Pengelly, A. R., Copur, O., Jackle, H., Herzig, A. & Muller, J. A histone mutant reproduces the phenotype caused by loss of histone-modifying factor Polycomb. Science 339, 698–699, doi:10.1126/science.1231382 (2013).

54. Morgan, M. A. J. & Shilatifard, A. Reevaluating the roles of histone-modifying enzymes and their associated chromatin modifications in transcriptional regulation. Nat Genet 52, 1271–1281, doi:10.1038/s41588-020-00736-4 (2020).

55. Zhang, T., Zhang, Z., Dong, Q., Xiong, J. & Zhu, B. Histone H3K27 acetylation is dispensable for enhancer activity in mouse embryonic stem cells. Genome Biol 21, 45, doi:10.1186/s13059-020-01957-w (2020).

56. Leatham-Jensen, M. et al. Lysine 27 of replication-independent histone H3.3 is required for Polycomb target gene silencing but not for gene activation. PLoS Genet 15, e1007932, doi:10.1371/journal.pgen.1007932 (2019).

57. Sankar, A. et al. Histone editing elucidates the functional roles of H3K27 methylation and acetylation in mammals. Nat Genet, doi:10.1038/s41588-022-01091-2 (2022).

58. Csordas, A. On the biological role of histone acetylation. Biochem J 265, 23–38, doi:10.1042/bj2650023 (1990).

59. Suka, N., Suka, Y., Carmen, A. A., Wu, J. & Grunstein, M. Highly specific antibodies determine histone acetylation site usage in yeast heterochromatin and euchromatin. Mol Cell 8, 473–479, doi:10.1016/s1097-2765(01)00301-x (2001).

60. Ruiz, P. D. & Gamble, M. J. MacroH2A1 chromatin specification requires its docking domain and acetylation of H2B lysine 20. Nat Commun 9, 5143, doi:10.1038/s41467-018-07189-8 (2018).

61. van der Velde, A. et al. Annotation of chromatin states in 66 complete mouse epigenomes during development. Commun Biol 4, 239, doi:10.1038/s42003-021-01756-4 (2021).

62. Luo, Y. et al. New developments on the Encyclopedia of DNA Elements (ENCODE) data portal. Nucleic Acids Res 48, D882–D889, doi:10.1093/nar/gkz1062 (2020).

63. Khan, A. & Zhang, X. dbSUPER: a database of super-enhancers in mouse and human genome. Nucleic Acids Res 44, D164–171, doi:10.1093/nar/gkv1002 (2016).

64. Ying, Q. L., Nichols, J., Evans, E. P. & Smith, A. G. Changing potency by spontaneous fusion. Nature 416, 545–548, doi:10.1038/nature729 (2002).

65. Li, H. & Durbin, R. Fast and accurate short read alignment with Burrows-Wheeler transform. Bioinformatics 25, 1754–1760, doi:10.1093/bioinformatics/btp324 (2009).

66. Li, H. et al. The Sequence Alignment/Map format and SAMtools. Bioinformatics 25, 2078–2079, doi:10.1093/bioinformatics/btp352 (2009).

67. Love, M. I., Huber, W. & Anders, S. Moderated estimation of fold change and dispersion for RNA-seq data with DESeq2. Genome Biol 15, 550, doi:10.1186/s13059-014-0550-8 (2014).

68. Frankish, A. et al. GENCODE reference annotation for the human and mouse genomes. Nucleic Acids Res 47, D766–D773, doi:10.1093/nar/gky955 (2019).

69. Quinlan, A. R. & Hall, I. M. BEDTools: a flexible suite of utilities for comparing genomic features. Bioinformatics 26, 841–842, doi:10.1093/bioinformatics/btq033 (2010).

70. Ramirez, F. et al. deepTools2: a next generation web server for deep-sequencing data analysis. Nucleic Acids Res 44, W160–165, doi:10.1093/nar/gkw257 (2016).

71. Yu, G., Wang, L. G. & He, Q. Y. ChIPseeker: an R/Bioconductor package for ChIP peak annotation, comparison and visualization. Bioinformatics 31, 2382–2383, doi:10.1093/bioinformatics/btv145 (2015).

72. Kent, W. J. et al. The human genome browser at UCSC. Genome Res 12, 996–1006, doi:10.1101/gr.229102 (2002).

73. Anders, S., Pyl, P. T. & Huber, W. HTSeq--a Python framework to work with high-throughput sequencing data. Bioinformatics 31, 166–169, doi:10.1093/bioinformatics/btu638 (2015).

74. Lawrence, M. et al. Software for computing and annotating genomic ranges. PLoS Comput Biol 9, e1003118, doi:10.1371/journal.pcbi.1003118 (2013).

75. Grau, J., Grosse, I. & Keilwagen, J.s PRROC: computing and visualizing precision-recall and receiver operating characteristic curves in R. Bioinformatics 31, 2595–2597, doi:10.1093/bioinformatics/btv153 (2015).

76. Durand, N. C. et al. Juicer Provides a One-Click System for Analyzing Loop-Resolution Hi-C Experiments. Cell Syst 3, 95–98, doi:10.1016/j.cels.2016.07.002 (2016).

